# A system-wide quantitative map of RNA and protein subcellular localisation dynamics

**DOI:** 10.1101/2022.01.24.477541

**Authors:** Eneko Villanueva, Tom Smith, Mariavittoria Pizzinga, Mohamed Elzek, Rayner M. L. Queiroz, Robert F. Harvey, Lisa M Breckels, Oliver M. Crook, Mie Monti, Veronica Dezi, Anne E. Willis, Kathryn S. Lilley

**Author notes:** These authors contributed equally and are in descending alphabetical order (Eneko Villanueva, Tom Smith, Mariavittoria Pizzinga). Correspondence: Kathryn Lilley or Anne Willis.

## Abstract

Existing methods to analyse RNA localisation are constrained to specific RNAs or subcellular niches, precluding the cell-wide mapping of RNA. We present Localisation of RNA (LoRNA), which maps, at once, RNAs to membranous (nucleus, ER and mitochondria) and membraneless compartments (cytosol, nucleolus and phase-separated granules). Simultaneous interrogation of all RNA locations allows the system-wide quantification of RNA proportional distribution and the comprehensive analysis of RNA subcellular dynamics. Moreover, we have re-engineered the LOPIT (Localisation Of Proteins by Isotope Tagging) method, enabling integration with LoRNA, to jointly map RNA and protein subcellular localisation. Applying this framework, we obtain a global re-localisation map for 31839 transcripts and 5314 proteins during the unfolded protein response, uncovering that ER-localised transcripts are more efficiently recruited to stress granules than cytosolic RNAs, and revealing eIF3d is key to sustain cytoskeletal function. Overall, we provide the most exhaustive map to date of RNA and protein subcellular dynamics.

## Introduction

Compartmentalisation of eukaryotic cells and the dynamic distribution of macromolecules such as RNA and protein across these compartments are vital for cell function. There are numerous different classes of RNA species that perform their function in a spatially restricted manner, for example by creating translation “hotspots” so that newly synthesised proteins can act at precise locations without disturbing cellular protein homeostasis^1^. The ability to map RNA species to different subcellular compartments, and to determine how they relocalise upon perturbation, is thus key to understanding cellular homeostasis^2,3^. Despite the role that localisation of the transcriptome plays in cell function, cell-wide methods to study RNA subcellular localisation are limited.

The localisation of a single RNA transcript can be determined using smFISH^4^, while the RNA content of specific organelles can be explored using proximity-dependent biotinylation techniques, such as APEX-seq^5,6^. As the application of APEX-seq only enables interrogation of a single subcellular compartment per experiment, it does not provide a cell-wide view of RNA localisation. Furthermore, combining multiple APEX-seq experiments to generate a more holistic cellular map is time consuming and is not compatible with measuring proportional quantification of RNAs at different localisations. Previous attempts to determine the localisation of RNA in a cell-wide manner have been restricted by technical limitations, including localisation-independent RNA clustering and length-dependent RNA-localisation biases^7,8^. Obtaining a comprehensive map of RNA localisation, therefore, remains an ongoing challenge.

A number of approaches have been developed to determine the subcellular localisation of proteins based on protein correlation profiling. These cell-wide methods involve cell fractionation, reordering of the subcellular content, and the use of localisation-specific abundance profiles of marker proteins to determine the subcellular localisation of the complete proteome^9–12^. One such technique is the Localisation Of Proteins by Isotope Tagging (LOPIT) method^13,14^. Since all compartments are mapped simultaneously, LOPIT is especially suited to study protein re-localisation in dynamic conditions^15^, providing a distinct advantage over other methodologies, where few proteins (immuno-fluorescence; IF) or specific locations (proximity-dependent biotinylation; BioID, APEX2) are measured per experiment. However, the LOPIT framework is not compatible with simultaneous mapping of RNA since it was developed solely for protein subcellular mapping.

Here, we have developed a new simple method to study cell-wide subcellular localisation of RNA (termed LoRNA). Importantly, LoRNA allows the proportional estimation of RNA species across multiple locations. LoRNA is based on the same concept as LOPIT, but with modified fractionation approaches to ensure no RNA distribution bias. To demonstrate its utility, we apply LoRNA to U-2 OS cells, to produce a comprehensive subcellular map of the transcriptome, uncovering the extent to which lncRNA is cytosolically distributed and features of RNA species that contribute to their subcellular distribution.

Additionally, the cell fractionation approaches we have developed for LoRNA are compatible with LOPIT, allowing simultaneous cell-wide maps of the proteome and transcriptome to be created. To demonstrate the use of this integrative framework in a dynamic context, we have simultaneously applied LoRNA and LOPIT to examine the redistribution of RNA and protein during the activation of the unfolded protein response (UPR). The UPR is triggered by the accumulation of unfolded proteins in the ER lumen^16^. Its activation reduces global protein synthesis rates, and this results in the loss of RNA targeting to the ER, the formation of stress granules (SGs), and the upregulation of stress-response genes^17^. Importantly, its deregulation is associated with disease states including neurodegeneration^18^, cancer progression^19^ and diabetes^20^.

Our results expose a major reorganisation of the transcriptome and proteome upon the UPR activation and reveal that RNA species associated with the ER are more readily recruited to SGs than cytosolic RNAs. Our data show that RNAs encoding cytoskeletal proteins are retained and targeted to the periphery of organelles during UPR and suggest this occurs through binding by eIF3d.

Altogether, applying this integrative approach generates the most comprehensive map of RNA and protein cell-wide localisation dynamics to date, which can be readily explored via the following dedicated open-source resource: https://proteome.shinyapps.io/density_lorna_rnaloc_gene/.

## Results

### Simultaneous RNA and protein subcellular localisation

Organelles can be distinguished by their physicochemical properties, and we have previously shown that this approach can be used to simultaneously sort all organelles and map protein subcellular localisation^14,21^. We reasoned that this principle could be applied to map the subcellular localisation of RNA in all niches at once. To demonstrate this, we developed a new method to sort the entire cellular content based on the density of its constituents and interrogate the localisation of RNA, which we refer to as LoRNA (Fig. 1a). Crucially, unlike previous methods^8^, the cell lysate is loaded into the density gradient at the density of free RNA (1.17 g/ml). Therefore, organelle-associated RNAs migrate upwards in the density gradient, while cytosolic RNA-protein complexes migrate downwards. This loading scheme avoids cytosolic RNA contamination of organelle-enriched fractions and allows RNA to reach equilibrium faster (Supplementary Fig. 1a). After equilibrium density centrifugation, three distinct bands were observed in the gradient, which were enriched in Endoplasmic Reticulum (ER) and mitochondria, nucleus, and cytosol proteins, respectively (Supplementary Fig. 1 b,c). Performing RNA-Seq along the gradient revealed distinct profiles resulting in separate clusters for RNAs known to localise to the mitochondria, ER, nucleolus, nucleus and cytosol (Fig. 1b,c Supplementary Fig. 1d, Supplementary table 1,2). To our knowledge, this is the first time the complete subcellular localisation of RNA has been successfully resolved at a cell-wide level. Furthermore, a novel sub-cytosolic RNA sedimentation profile was discovered by semi-supervised clustering, hitherto referred to as ‘cytosol light’, which we explore in more detail later (Fig. 1b, Supplementary Fig. 1e,f).

**Figure 1.**
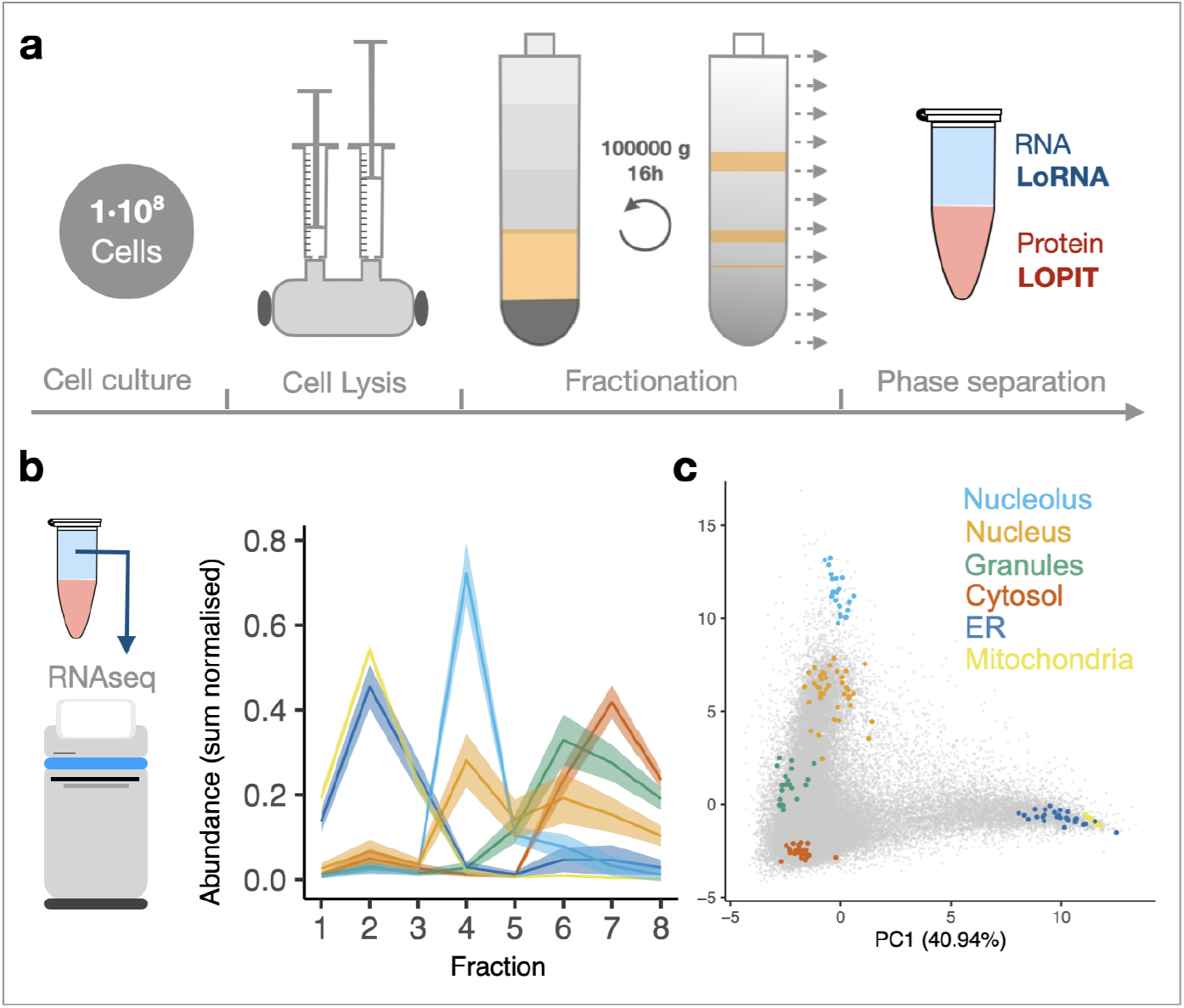
RNA subcellular localisation. **a,** Schematic representation of LoRNA. Cells are lysed and fractionated by density equilibrium centrifugation. RNA and protein are extracted from each fraction to perform LoRNA and LOPIT. Cell lysate and gradient banding pattern are represented in yellow. Aqueous and organic phases are blue and red respectively **b**, Application of LoRNA to U-2 OS cells. Mean profiles for RNA markers along pooled gradient fractions in a single experiment. Shaded regions denote +/- one standard error. **c,** PCA projection of RNA profiles across three replicate experiments, with marker RNAs highlighted.

Importantly, we engineered this new cell fractionation method to allow the simultaneous mapping of RNA and protein localisation (Fig 1a). To map protein subcellular localisation we adapted the LOPIT framework and quantified protein abundance in the same gradient fractions by mass spectrometry (Supplementary Fig 1 g,h). Analysis of protein mapping accuracy by classifying organelle marker proteins resulted in F1 scores (harmonic mean of precision and recall) of 0.71 - 1 for each organelle, demonstrating the high resolution of this novel density-based LOPIT (dLOPIT) method (Sup Fig. i, Supplementary table 3). These results demonstrate that the combination of LoRNA with dLOPIT allows the simultaneous mapping of the transcriptome and proteome subcellular localisation.

### Simultaneous localisation of RNA content permits its quantification at different subcellular compartments

To estimate the proportional content of each RNA in every localisation, relative RNA abundances were adjusted to account for the total RNA content per fraction. This allowed us to decompose the RNA profiles into the constituent contributions from each localisation, as determined from the profiles of RNAs with known localisations (Fig. 2a). In this way, we estimated the proportional localisation of 31839 transcript isoforms (13142 genes). Transcript-level RNA mapping retains valuable information on the differential localisation of splicing variants (Sup Fig 2a). We therefore use transcript-level proportions, unless specified otherwise. As expected, RNAs exist on two major axes: Nucleus:Cytosol and Cytosol:Membrane, with expected proportions obtained for RNAs known to be extremely enriched in specific locations (Fig. 2b).

**Figure 2.**
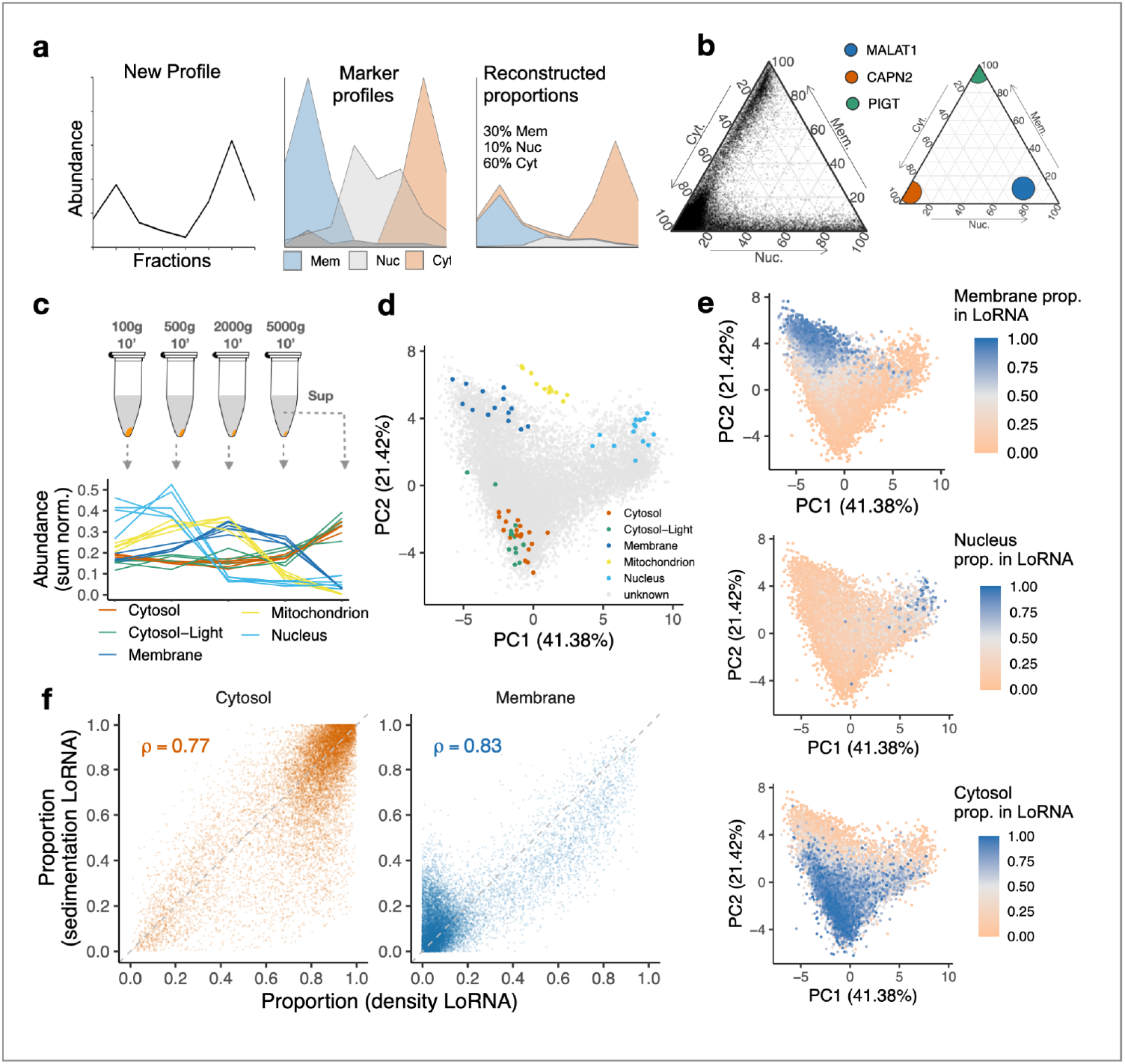
System-wide quantification of RNA localisation. **a,** Schematic representation of how localisation proportions are estimated. New profile (left) is decomposed into different proportions of each marker profile (middle) to approximate the original profile (right). **b**, Cytosol, membrane and nuclear (including nucleolus) proportions for all transcripts (left) and selected localisation-enriched transcripts (right). **c,** Schematic representation of cell fractionation by sedimentation coefficient and linear profiles of the localisation markers. **d**, PCA projections of RNA profiles in **c**, with markers highlighted. **e**, Membrane, nucleus and cytosol proportions obtained by equilibrium density centrifugation-based LoRNA projected on the sedimentation-based-RNA map. **f**, Correlations between proportions obtained using different fractionation approaches. Cytosol proportions by density is the sum of cytosol and cytosol-light proportions. Membrane proportion by sedimentation is the sum of mitochondrial and ER proportions.

To comprehensively validate our system-wide RNA localisation results, we developed an orthogonal method to sort the cellular content by a different physicochemical property, namely the sedimentation coefficient of the different organelles instead of density. Previous attempts to study RNA localisation using this concept applied high *g*-forces^7^, and analysis of the published data highlights that this creates an RNA-length dependent localisation bias (Supplementary Fig. 2a,b). We therefore optimised the fractionation speeds and times to avoid RNA-length-dependent sedimentation bias, whilst still separating the major subcellular localisations (Fig. 2c & Supplementary Fig. 2b,c,d, Supplementary table 1, 2). While the low g-force required to avoid this bias precludes sub-cytosolic resolution, differential centrifugation provides better separation between mitochondria and ER than density centrifugation (Fig. 2d). The projection of the densitybased RNA proportions onto the map obtained with our orthogonal approach showed a remarkable agreement for membranes, nucleus, and cytosol (Fig. 2e). In addition to the excellent separation of the main RNA localisation niches, reliable estimates of cytosol and membrane proportions were also achieved (Fig. 2f, Supplementary Fig 2e). These proportions show Pearson’s correlations of 0.77 and 0.83, respectively, confirming the reliability of our estimates. Thus, LoRNA provides an accurate quantitative system-wide map of RNA localisation, for the first time.

### Key RNA features drive subcellular localisation

We next explored the RNA features associated with proportional localisation. While mRNAs are observed throughout the cell, and relatively depleted from the nucleus, lncRNAs were observed to be strictly within the nucleus:cytosol axis (Fig. 3a left). Although lncRNAs are more nuclear localised than mRNAs, the overall distribution of lncRNAs was more cytosolic than we anticipated (Fig. 3b). Notably, comparing the proportions between our two cell fractionation approaches, we observed that 59.7% of lncRNAs were consistently cytosolic in both approaches. As expected, higher ribosome association correlates with greater cytosol localisation^22^ (Supplementary Fig. 3a). Interestingly, we found that cytosolic localisation can be predicted according to transcript length, presence of polyA tail and AU content, with short, polyadenylated and low AU content lncRNA being more cytosolic (Fig. 3c). This simple model is as accurate as a more complex model built from a penalised regression over a wide range of potential features (ROC AUC = 0.86 for both models; Supplementary Fig. 3b). We therefore favour this simpler model and suggest these are the key features driving cytosolic localisation for lncRNAs (Fig. 3c).

**Figure 3.**
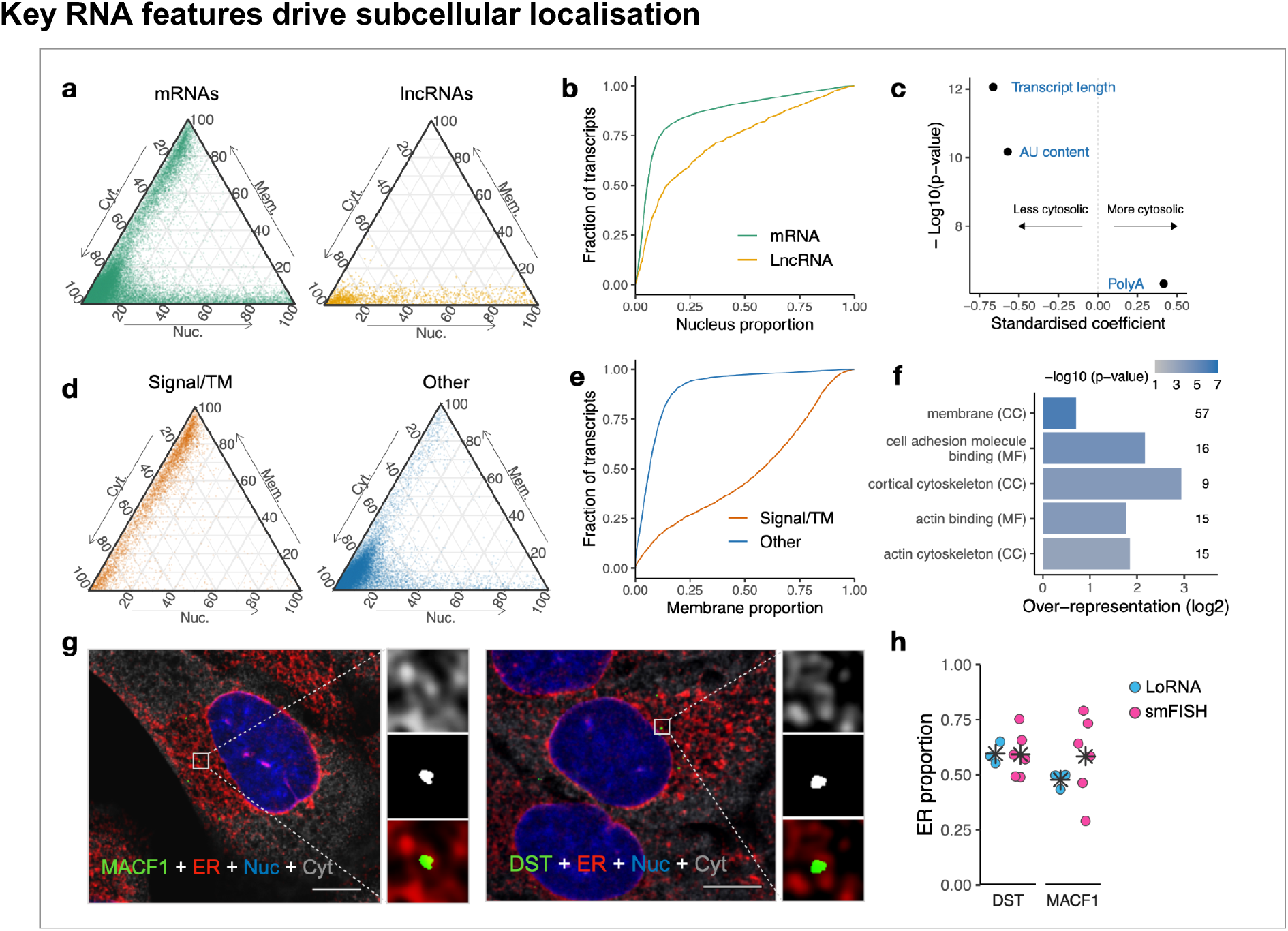
Features driving RNA localisation. **a,** Cytosol, nucleus and membrane proportions for mRNAs and lncRNAs. **b**, Empirical cumulative frequency distributions for nucleus proportions for mRNAs and lncRNAs. **c**, Coefficient estimates and p-values for logistic regression model of lncRNA cytosol proportions. **d**, Proportions for mRNAs encoding signal peptides and/or transmembrane (TM) domains. **e**, Membrane proportions for mRNAs shown in d. **f**, GO-terms significantly enriched in mRNAs not encoding signal peptides and/or TM domains but over 35% membrane localised. **g**, smFISH validation of ER localisation of MACF1 and DST RNAs not encoding signal peptides and TM domains. **h**, ER proportion in smFISH and LoRNA.

While generally we found mRNAs predominantly in the cytosol, RNAs encoding proteins with signal peptides and/or transmembrane domains (TMs) were much more prominently membrane localised, consistently with their co-translational targeting to the ER (Fig. 3d). Furthermore, our data confirmed the relationship between the distance from the first signal peptide / TM domain to the stop codon and the membrane proportion (Supplementary Fig. 3c)^23,24^. Surprisingly, we also identified 100 membrane-localised RNAs that did not encode a signal peptide or TM. These included the mRNA encoding the ER-localised signal recognition receptor subunit beta, which co-translationally binds to the alpha subunit^25^, and mRNAs encoding for mitochondrial proteins (Supplementary Fig. 3d), but most of the RNAs were not previously described as membrane localised. GO analysis revealed a significant enrichment for terms associated with membrane or cytoskeleton localisation (Fig. 3f), suggesting potential localised translation at the surface of the membranes. smFISH quantification of the ER association for the Microtubule-actin cross-linking factor 1 (MACF1) and Dystonin (DST) mRNAs confirms the localisation of transcripts encoding cytoskeletal proteins at the ER (Fig 3 g,h).

### Global assessment of transcriptome and proteome relocalisation

We next combined LoRNA and dLOPIT to interrogate RNA and protein re-localisation upon activation of the unfolded protein response. The UPR is an adaptive signalling pathway induced by the accumulation of unfolded proteins in the ER lumen to reinstate homeostasis by reducing the protein folding load and increasing the protein folding capacity of the ER. To induce the UPR, we inhibited the SERCA Ca^2+^ pump by treating U-2 OS cells with 250nM of Thapsigargin (TG) for 1 h. This reduces calcium in the ER lumen, impairing protein folding and activating the UPR, which, in turn, rapidly induces a global translational shutdown through phosphorylation of eIF2α (Supplementary Fig. 4a,b). This results in the formation of stress granules (SGs) (Supplementary Fig. 4c) and the increased expression of UPR genes like XBP1 and CHOP (Supplementary Fig. 4d).

LoRNA allows quantification of the redistribution of RNA upon inhibition of translation, with RNA migrating from the membranes towards the cytosol (Fig. 4a, Supplementary Fig. 4e). Surprisingly, we found a pronounced relocalisation of RNAs to the cytosol light, both from the cytosol and from membranes (Fig. 4b). dLOPIT analysis of protein re-localisation identified 73 proteins that were differentially localised 1h post-stimulation (Fig. 4c,d). The Golgi apparatus was the most affected organelle after the short stimulation, as expected given its role in the trafficking of newly synthesised proteins. Interestingly, six proteins lost their co-localisation with the ribosome upon UPR, including proteins known to relocalise to stress granules, such as STAU2 and PABPC1. The linear profiles for these proteins along the gradient fractions show that 4/6 increase in abundance in the fraction we previously observed to differentiate the cytosol light profile for RNAs (Fig. 4e, Supplementary Fig 4f). We therefore hypothesised that the cytosol light profile may represent RNP granules. To test this, we identified further proteins whose profile overlapped the cytosol light RNA profile, using the profile of these 4 proteins in UPR. We identified 22 proteins with a matching UPR profile, including 5 members of the P-body associated CCR4-NOT complex^26^, P-body proteins DCP1A/B^27^ and stress granule components YBX3^28^, SECISBP2^28^ and CASC3^29^. Furthermore, the profile of GFP-tagged DCP2 and G3BP1 proteins in the density gradients confirmed that cytosolic phase-separated particles sediment at the density ranges of the cytosol light (Supplementary Fig. 4g,h, supplementary methods), suggesting that the cytosol light RNA profile represents the distribution of phase-separated cytosolic RNP particles. Thus, the integration of LoRNA and dLOPIT allows, for the first time, the simultaneous quantification of the transcriptome and proteome re-localisation in membrane enclosed organelles and membraneless compartments.

**Figure 4.**
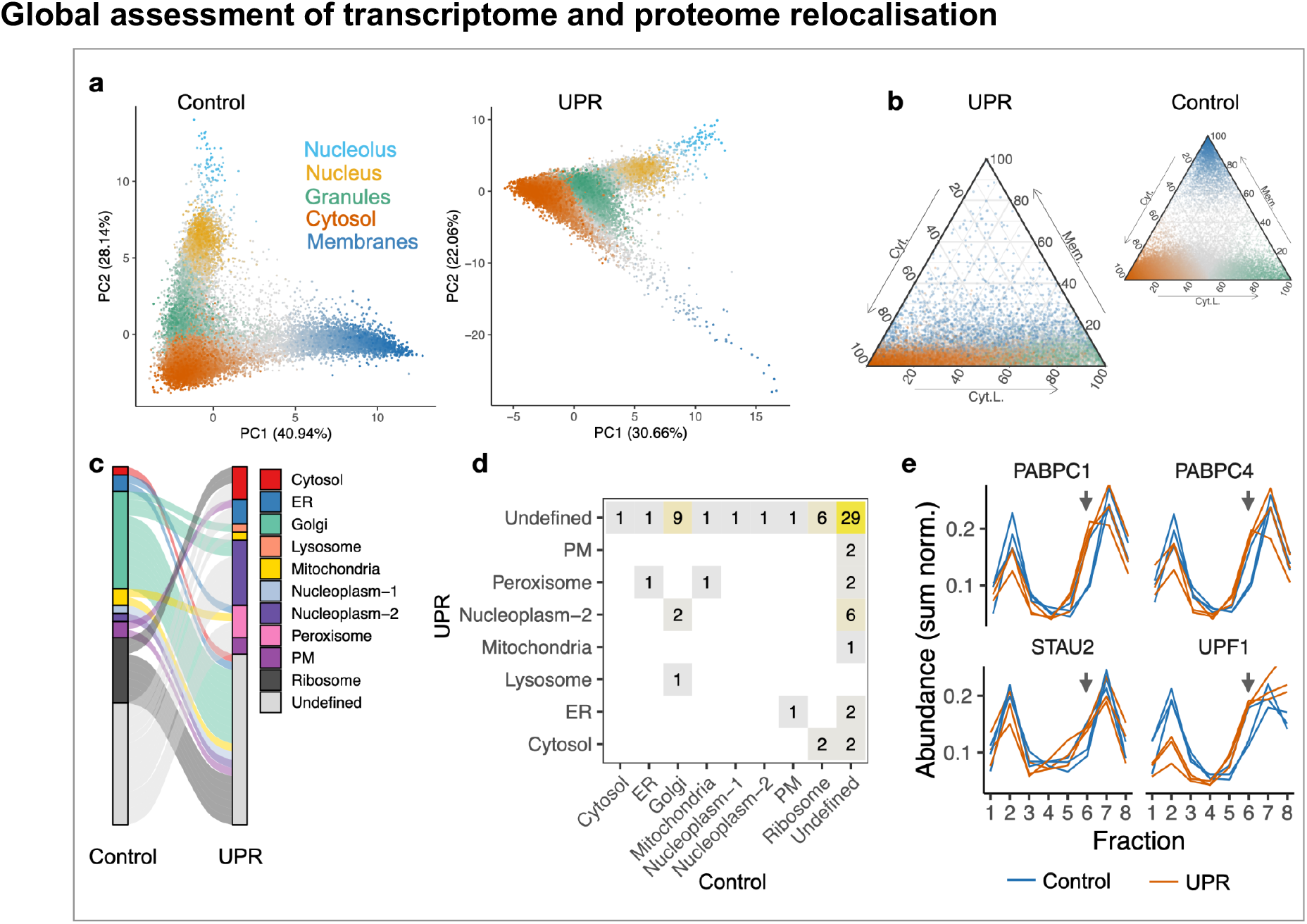
Transcriptome and proteome subcellular redistribution upon UPR. **a,** PCA projection of RNA profiles in control and UPR. Colour and intensity indicates RNA proportion for the primary localisation. PC2 in UPR is inverted. **b,** Cytosol, membrane and cytosol light proportions in control and UPR. Point colour indicates RNA major localisation (excluding nucleus and nucleolus) in control, with colour intensity denoting proportion. **c-d**, Differential protein localisation in control and UPR. **e**, Abundance profiles for stress granule proteins relocalising from ribosomes to the fraction discriminating the cytosol-light-RNA profile under UPR, grey arrows mark cytosol light discriminating fraction.

### RNA features driving granule localisation

As the proteins relocalising to the cytosol light upon UPR are associated with RNA condensation into phase-excluded particles, we further characterised the RNA composition of this fraction and interrogated the features of the enriched RNAs. RNAs with higher cytosol light abundance were correlated with lower ribosome association and longer transcript length (Fig. 5a,b), both features associated with RNAs partitioning to granules^30,31^. Furthermore, RNAs enriched in P-bodies using fluorescence-activated particle sorting have a similar distribution to the cytosol light profile^32^ (Supplementary Fig. 5a). Altogether, this provides strong evidence that our cytosol light profiles represent granule localisation and we thus refer to this localisation accordingly henceforth.

**Figure 5.**
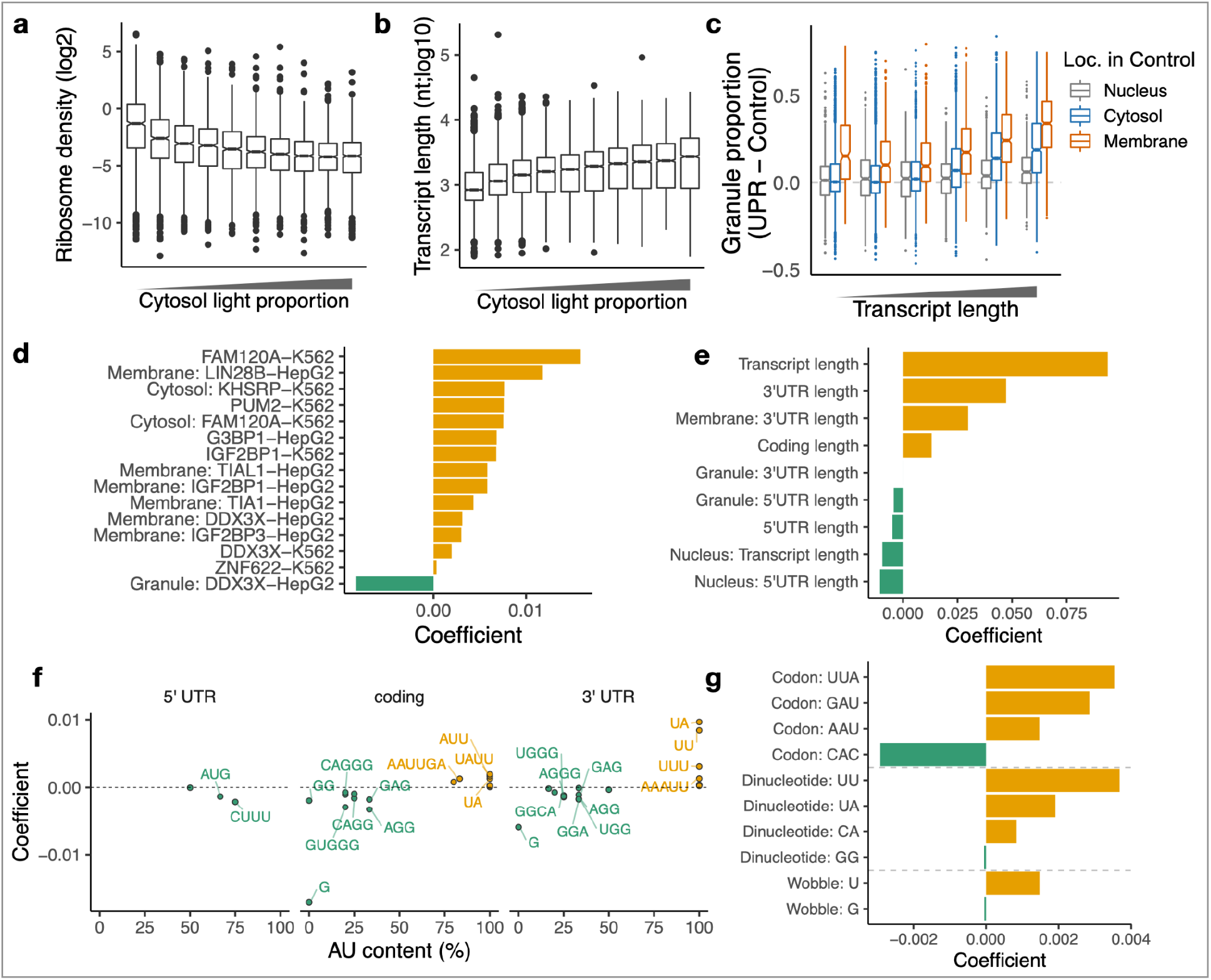
Analysis of the characteristics driving RNAs to granules. **a-b**, Ribosome density^33^ and transcript length for RNAs split into deciles of cytosol light proportion, shown as Tukey boxplots. **c**, Relocalisation to granules (cytosol light) in UPR. RNAs are binned by transcript length and split by localisation in control conditions. **d**, Coefficients for features selected by lasso regression model of relocalisation to granules. Features describe RBP binding or the interaction between RBP binding and localisation in control. RBP eCLIP cell line indicated in feature name. Positive coefficients mean increased relocalisation to granules. **e**, Coefficients for RNA length features. **f**, Coefficients and AU content for kmer features. **g**, Coefficients for codon features.

Our quantification of RNA recruitment to granules following induction of the UPR agrees with a recently published targeted study^34^ (Supplementary Fig. 5b). Importantly, LoRNA provides cell-wide information on RNA localisation prior and post stimulation. This allowed us to uncover that, although many cytosolic RNAs migrate to granules upon UPR, membrane RNAs migrating to granules do so at a higher proportion (Fig. 5e, Supplementary Fig. 5c). To fully characterise the features driving RNAs towards granules upon UPR activation, we modelled the contribution of a broad range of RNA features, including transcript length, localisation prior to stress, RBP binding, kmer content, codon usage, and presence of IRES or upstream ORFs. Overall transcript length, especially longer 3’UTRs in membrane RNAs, was observed to be the greatest positive predictor of relocalisation to granules (Fig. 5f). RBPs whose binding was positively predictive of RNA relocalisation to granules included the granule proteins FAM120A, IGF2BP1, and G3BP1 (Fig. 5g) with binding of canonical stress granule proteins TAIL1, TIA1 and IGFBP3 to membrane-localised RNAs also showing a significant predictive value. Intriguingly, unexpected associations between RBP binding and relocalisation to granules were observed, including LIN28B in membrane-localised RNAs and ZNF622, which is involved in ribosome subunit joining^35^ and upregulated upon viral infection and UPR stress^36^. Importantly, we found that ZNF622 relocalises from the cytosol to the ER upon UPR (Supplementary Fig. 5d,e), a step that may be required for the membrane-RNA recruitment to granules.

A higher AU content in the coding and 3’UTR regions was predictive of greater relocalisation to granules (Fig. 5f). The latter fits with the known role of 3’UTR AU-rich elements (AREs) in regulating RNA stability. Interestingly, higher G content, but not C content, in the coding region was a predictor of lower granule relocalisation (Fig. 5f). In addition, the frequency of AUG kmers in the 5’UTR was predictive of lower relocalisation, although presence of annotated uORFs was not. This may indicate that the presence of an AUG at the 5’UTR inhibits granule association, regardless of the ORF capacity of the sequence. Finally, while overall codon optimality was not a selected predictive feature, the frequency of UUA, GAU, AAU, and CAC codons were predictive of granule relocalisation (Fig. 5g). These codons include 3 of the 4 codons that require queuosine-modified tRNAs for their decoding (p=0.003, Fisher’s exact test), pointing to a potential role of this modification in the stress response and RNA recruitment to phase-separated particles.

### RNAs coding for cytoskeletal proteins retain organelle association upon UPR

Despite the global loss of RNA from the membranes, some RNAs remain membrane-enriched upon UPR (Fig. 6a). We modelled this relationship using a generalised additive model and identified RNAs with unexpectedly high membrane localisation after UPR. Notably, RNAs with longer sequences between the signal sequence and stop codon are retained more efficiently in membranes, suggesting the involvement of the signal recognition particle ^23^ and therefore active reassociation with the membranes (Fig. 6b). To gain a deeper understanding on the mechanisms of RNA retention in the periphery of organelles upon UPR activation, we then identified RBPs previously found to interact with these RNAs using publicly available eCLIP data ^38^ which could explain this higher membrane localisation. We observed that eIF3d binding was associated with the greatest degree of membrane localisation (Fig. 6c, Supplementary Fig. 6a). We further confirmed this using complementary data from a targeted CLIP experiment that specifically identified RNAs bound by eIF3d at their 5’cap^37^ (Fig. 6d). Comparing the membrane-localised RNAs bound by eIF3d to those not bound to it, we observed an over-representation for GO terms relating to the actin cytoskeleton, cell morphogenesis, and focal adhesions (Fig. 6e). We therefore postulate that eIF3d could be required for remodelling the cytoskeleton in processes that may be dependent on localised translation like cell migration, particularly during activation of the UPR. Indeed, knocking down eIF3d (Supplementary Fig. 6b) reduced cell migration overall, with a significatively stronger effect in cells undergoing UPR (Fig. 6f,g, Supplementary Fig. 6c). eIF3d has been previously shown to promote recruitment of specific mRNAs to ribosomes during ER stress^39^. Our results suggest that eIF3D is required for continued translation of actin cytoskeleton components localised in the periphery of the membranes during UPR.

**Figure 6.**
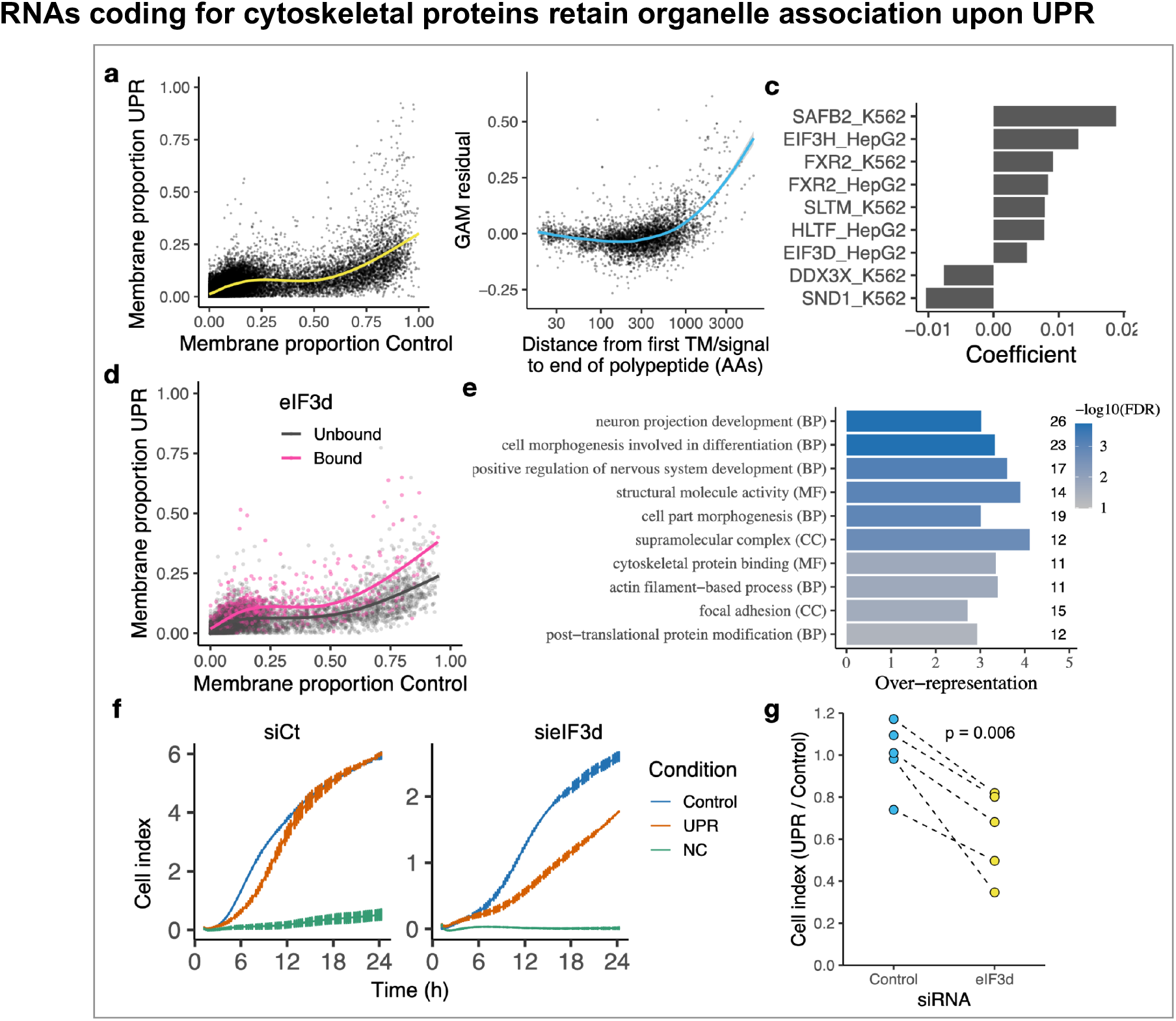
Analysis of the RNAs retaining membrane association under UPR. **a,** Membrane proportions in control and UPR. Yellow line indicates fit from generalised additive model (GAM) with cubic regression spline. **b**, Relationship between residual from GAM and the distance between the first signal or TM domain and stop codon. Blue line indicates smoothed fit by LOESS local regression. **c**, Coefficients for RBPs which are predictors of GAM residuals for RNAs which do not encode a signal peptide/TM domain. RBP eCLIP cell line indicated in feature name. **d**, Membrane proportions as per a, for RNAs bound by eIF3d according to eCLIP or Subunit-Seq^37^. **e**, GO terms enriched in membrane RNAs bound by eIF3d, relative to all membrane RNAs. **f**, Representative U-2 OS cell migration time series for eIF3d knock-down and siRNA Control (siCt) cells in control and UPR conditions. **g**, Quantification of cell migration at 24 h post UPR induction in eIF3d knock-down and siRNA Control cells (n=5).

## Discussion

In this work, we have developed an integrative framework coupling two new methods, LoRNA and dLOPIT, to generate the first cell-wide simultaneous map of RNA and protein subcellular localisation in membranous (nucleolus, ER, and mitochondria) and membraneless (cytosol, nucleolus, and cytosolic granules) compartments. By precisely reconstructing RNA localisation, LoRNA allows quantitative determination of the complete subcellular distribution of each RNA. Coupling LoRNA and dLOPIT, we characterise the dynamic transcriptome and proteome subcellular redistribution upon UPR. We establish the RNA features driving relocalisation to granules, identify transcripts that are targeted to the periphery of the organelles during the unfolded protein response, and suggest a role for eIF3d in maintaining cytoskeleton function upon UPR activation.

When interpreting LoRNA proportions, it is important to consider that the RNA marker profiles that are used to estimate localisation proportions do not represent spatial coordinates in the cell, but rather the typical profile for RNAs which predominantly reside in that localisation. For example, the distribution of the cytosolic RNA markers include nascent RNA copies that are nuclear-localised. As such, proportions are with respect to RNAs that are paradigmatic representatives of localisation, and do not represent absolute localisations. This subtle distinction is important when comparing LoRNA proportions to orthogonal methods such as smFISH where the localisation of individual molecules is examined. Furthermore, in cases where gradients are of interest, such as across the cytosol of a polarised cell, targeted methods such as FISH or IF are required as any high-throughput method involving cell lysis will necessarily result in a loss of subcellular coordinates. When RNA compartments not amenable to biochemical fractionation (e.g. the nucleopore) are of interest, alternative methods like APEX-Seq may be applicable. Nevertheless, multiple APEX-based experiments need to be combined to cell-wide map RNA localisation, making this approach incompatible with the proportional estimation of RNA localisation and specially challenging to apply in dynamic systems or to study multiple biological systems. Nonetheless, APEX and LoRNA can function as complementary approaches to interrogate molecular subcellular distribution at different resolutions.

One of our most striking findings regarding RNA relocalisation after UPR activation is that membrane-localised RNAs are recruited more efficiently to granules than cytosolic RNAs. This conflicts with transcriptomic studies that have claimed ER-targeted RNA are depleted from SGs^40^. However, targeted methods to purify SGs typically focus on specific engineered bait proteins and require multiple purification steps, while LoRNA recovers granules regardless of their specific composition, based on their distinct density sedimentation profile, which may explain the apparent discrepancy between SG transcriptomics and LoRNA. Since our results suggest ER-RNA relocalisation to granules during stress is widespread, we speculate that novel subtypes of stress-induced granules may be the ultimate destination for membrane RNAs upon UPR activation, opening new avenues for future research. Furthermore, we observed previously uncharacterised predictors of RNA relocalisation to granules. Of particular note, we found that ZNF622 relocalises to the ER upon UPR and RNAs containing ZNF622 binding sites relocalise more readily to granules. Given that ZNF622 has been recently shown to inhibit ribosome subunit joining^35^, and may be involved in the ER associated degradation (ERAD) machinery or ER quality control following viral infection^36^, ZNF622 relocalisation to the ER may be required to inhibit ER-localised translation and drive ER-targeted RNAs to granules upon UPR. Additionally, the relocalisation of RNA to granules is associated with altered frequencies for 3 codons that require queuosine-modification in the U-wobble position of the cognate tRNA. Considering the emerging role that tRNA modifications play in translation^41,42^, it would be interesting to explore if a tRNA-specific co-translational mechanism could regulate the recruitment of RNAs to SGs.

The precise measurement of RNA proportionality afforded by LoRNA allowed characterisation of the relocalisation of RNA away from membranes in unprecedented detail. Intriguingly, we found that despite the general loss of mRNA from the ER upon UPR^43^, many RNAs are partially retained in association with organelles, or even migrate to the ER. Notably, we found that many of these mRNAs encode for proteins involved in cytoskeletal remodelling. Furthermore, eIF3d binding correlates with membrane localisation and knocking it down impairs cell migration, a process highly dependent on cytoskeletal remodelling. Actin cytoskeleton remodelling plays a key role to restore Ca^2+^ homeostasis in the ER upon UPR through promotion of ER-PM contact^44^. Separately, eIF3d has been found to maintain RNA translation upon stress^39^. Altogether, we speculate that the cytoskeleton remodelling required to overcome UPR is eIF3d-dependent and that it involves the in-situ translation of cytoskeletal mRNAs.

In summary, our work provides the first quantitative system-wide determination of RNA localisation and re-localisation during the UPR. The resulting data has invaluable potential for future studies characterising the functions of specific RNAs and proteins. To facilitate this, we have generated a user-friendly graphical interface to explore our data available at https://proteome.shinyapps.io/density_lorna_rnaloc_gene/. LoRNA allows for the unbiased cell-wide determination of RNA compartmentalisation. This will support a much-needed paradigm shift from the study of RNA localisation through relative enrichments between two localisations, towards a cell-wide analysis of RNA proportional distribution. We anticipate others will build upon LoRNA. For example, coupling it with direct RNA sequencing methods would enable the precise interrogation of the role of RNA modifications in RNA localisation. By enabling simultaneous characterisation of proteome and transcriptome relocalisation, the combination of LoRNA with dLOPIT represents a transformative approach to study how the cell coordinately responds to physiological signals, stress conditions, exogenous cues, or infectious pathogens. Importantly, integrative research combining LoRNA and dLOPIT can contribute to the translation of molecular biology observations to medical benefits by fostering new studies into the role of RNA and protein localisation dynamics in cell homeostasis and disease.

## Methods

### Cell culture and UPR induction

U-2 OS cells were obtained from the American Type Culture Collection (ATCC), maintained in McCoy’s A5 media (Gibco-BRL) supplemented with 10% of FBS (Gibco-BRL), at 37 °C and 5% CO2, and regularly tested for mycoplasma contamination with negative results. UPR was induced by directly adding 250 nM of Thapsigargin (TG, UPR) or equivalent volume of DMSO (control) to cells at 90% confluency. Cells were incubated with TG or DMSO for 1 h at 37 °C unless specified otherwise.

### Density based cell fractionation

7ml discontinuous density gradients of 15, 25 and 25% iodixanol (Optiprep, Stemcell technologies), 0.25 M sucrose, 75 mM KCl, 5 mM MgCl_2_, 50 nM CaCl_2_, 10 mM HEPES pH 7.4 and EDTA free protease inhibitor were prepared in polyallomer optiSeal ultracentrifuge tubes (11.2 ml capacity; Beckman Coulter) gradients were allowed to diffuse 1h at 20 °C. Partially diffused gradients were stored at 4°C for 1 h while cells were prepared for fractionation. Cells were cultured in 500 mm^2^ pates until 90% confluence, using a single plate per replica per condition. Cells were treated with TG or DMSO for 1 h. After treatment, cells were washed twice with PBS and detached using EDTA-free trypsin (Thermo Scientific) for 10 min. Trypsin was quenched by adding equal volume of media (supplemented with DMSO or TG). Detached cells were transferred to a 50 ml tube and spun down 10 min at 250 g. Cell pellets were washed twice with ice cold PBS and resuspended in 900 μl of lysis buffer (0.25 M sucrose, 75 mM KCl, 5 mM MgCl_2_, 50 nM CaCl_2_, 10 mM HEPES pH 7.4 and EDTA-free protease inhibitor) and lysed with a ball bearing homogenizer (Isobiotec) on ice. 50 μl of lysate was stored at −80°C as total cell lysate. Cell lysate iodixanol and ion concentration was adjusted by adding 1.5 ml of 50% iodixanol solution (in 75 mM KCl, 5 mM MgCl_2_, 50 nM CaCl_2_, 10 mM HEPES pH 7.4 and EDTA free protease inhibitor) to a 1 ml cell lysate, and underlaid in the previously prepared density gradient with a 2.5 ml syringe and a wide-bore blunt-end needle (Sigma-Aldrich). Finally, a 40% iodixanol (in 75 mM KCl, 5 mM MgCl_2_, 50 nM CaCl_2_, 10 mM HEPES pH 7.4 and EDTA-free protease inhibitor) cushion was underlied until the tube was filled. Density gradients were centrifuged in a NVT65 fixed-angle near-vertical ultracentrifuge rotor (Beckman Coulter) in a Optima L-80 XP ultracentrifuge (Beckman Coulter) for 16 h at 100.000 g and collected using an auto Densi-Flow peristaltic pump fraction collector with a meniscus-tracking probe (Labconco) to obtain 20 fractions of 500 μl each. The refractive index (RI) of each fraction is measured with a Hand-held refractometer (Reichert) and the iodixanol concentration was calculated as Iodixanol% = (RI / 0.83) - 10.111, and fraction density calculated by d = m / V. The iodixanol concentration per fraction was adjusted to 30% in a volume of 600 μl. All fractions were frozen and dried by sublimation using Vacuum centrifuge with cold trap (Labconco, Refrigerated CentriVap concentrator). Dried pellets were solubilised in 1 ml of trizol (Thermo Scientific) and stored at −80 °C.

### Differential sedimentation speed based cell fractionation

Cells were cultured in 500 mm^2^ plates until 90% confluence, using a single plate per replicate. Five replicates were performed for the differential sedimentation speed based cell fractionation experiment. Cells were washed twice with PBS and detached using trypsin/EDTA (0.05%) (Thermo Scientific) for 5 min. Trypsin was quenched with an equal volume of media. Detached cells were transferred to a 50 ml falcon tube and spun down for 5 min at 200 g at 4 °C. The pellets were washed twice with ice cold PBS and resuspended in 1 ml of lysis buffer (75 mM KCl, 5 mM MgCl_2_, 50 nM CaCl_2_, 10 mM HEPES pH 7.4 and EDTA-free protease inhibitor) and homogenised on ice using ball bearing homogenizer (Isobiotec). A total lysate sample of 75 μl was obtained and stored at −80 °C. The remaining sample was fractionated into five consecutive fractions at centrifugation speeds (100, 500, 2000, and 5000 g) using the supernatant of every centrifugation as starting material for the next, with a Eppendorf Centrifuge 5424R. The supernatant of the last centrifugation was retained as the final fraction.

### RNA and protein sample precipitation

RNA and protein were obtained from trizol solubilised fractions by adding 200 μl of chloroform and phase partitioning the sample for 15 min at 12000 g at 4 °C. RNA was purified by collecting and transferring the trizol/chloroform upper aqueous phase to a new tube and the RNA precipitated with 750 μl of isopropanol (Sigma Aldrich) for 10 min at 16000 g. RNA pellets were washed twice with 70% ethanol and solubilised in 200 μl of RNAse-free water (Thermo Scientific). RNA samples were treated with DNAse in RNeasy columns (Qiagen) according to manufacturer instructions using the RNase-free DNAse Set kit (Qiagen). RNA concentration was measured with a DS-11 UV Spectrophotometer (Denovix). RNA samples were pooled as indicated in the supplementary table 4. Protein was purified by precipitating the trizol/chloroform lower organic phase (and interface) using 9:1 v:v of methanol:sample. Samples were solubilised in 1% SDS (Thermo Fisher Scientific) 100 mM TEAB (Sigma-Aldrich) using a Bioruptor sonicating bath (Diagenode). Protein concentration was measured with Pierce BCA Protein concentration assay kit (Thermo Fisher Scientific) on a spectrophotometer plate reader (Molecular Devices, SpectroMax M2).

### RNA sequencing

RNA sequencing libraries for the density fractionation were generated using 1 μg of RNA as starting material and depleting ribosomal RNA using Ribocop V3 (Lexogen). Post ribosomal RNA depletion, RNA content was measured with a Bioanalyzer pico kit (Agilent). 1% RNA SIRVs were spiked-in (Lexogen), and RNA seq libraries were generated and amplified using the CORALL RNA sequencing kit (Lexogen) according to the manufacturer instructions. All CORALL generated libraries were sequenced in parallel on 4 Novaseq S4 lanes (Illumina). RNA sequencing libraries for the differential sedimentation speed based cell fractionation experiment were performed using QuantSeq 3’ mRNA-Seq kit (Lexogen) according to manufacturer instructions. A total of 400 ng of total RNA and 0.1% RNA SIRVs (Lexogen) were used for library preparation of each fraction. All libraries were balanced, multiplexed and pooled, and run on two single-end Novaseq SP lanes (Illumina).

### Proteomic sample preparation

Samples were resuspended in 100 μL of 100 mM TEAB (Sigma-Aldrich), reduced with 10 mM DTT (Sigma-Aldrich) at room temperature for 60 min and alkylated with 40 mM iodoacetamide (Sigma-Aldrich) at room temperature in the dark for 60 min. Samples were digested overnight at 37 °C with 1 μg of trypsin (Promega). Subsequently, 1 μg of modified trypsin (Promega) was added, and the samples were incubated for 3-4 h at 37 °C. Samples were then acidified with TFA (0.5% (v/v) final concentration (Sigma-Aldrich) and centrifuged at 21,000 g for 10 min. The supernatant was immediately desalted.

For peptide clean-up and quantification, 200 μL of Poros Oligo R3 (Thermo Fisher Scientific) resin slurry (approximately 150-200 μL resin) was packed into Pierce centrifuge columns (Thermo Fisher Scientific) and equilibrated with 0.1% TFA. Samples were loaded, washed twice with 200 μL 0.1% TFA and eluted with 300 μL 70% acetonitrile (ACN) (adapted from^45^). From each elution, 10 μL was taken for Qubit protein assay (Thermo Fisher Scientific) quantification and the remaining sample was retained for MS.

TMT-10plex or TMTpro-16plex (Thermo Fisher Scientific) labelling from desalted peptides was performed according to the manufacturer’s protocol. Equal amounts of desalted peptides were labelled immediately after being quantified with Qubit protein assay (Thermo Fisher Scientific). Multiplexed TMT samples were then fractionated using high-pH reverse phase chromatography. In detail, the TMT-labelled peptide samples were resuspended in 100 μL of 20 mM ammonium formate pH 10 (Buffer A). The total volume of each sample was injected onto an Acquity UPLC BEH C18 column (2.1-mm i.d. × 150-mm; 1.7-μm particle size) on an Acquity UPLC System with a diode array detector (Waters) and the peptides were eluted from the column using a linear gradient of 4–60% (v/v) acetonitrile in 20 mM ammonium formate pH 10 over 50 min and at a 0.244 mL/min flow rate (with a total run time of 75 min). The gradient was set up as follows: 0 min-95% Buffer A–5% Buffer B (20 mM ammonium formate pH 10 + 80% (v/v) acetonitrile), 10 min-95% Buffer A–5% Buffer B, 60 min–25% Buffer A–75% Buffer B, 62min–0% Buffer A–100% Buffer B, 67.5min–0% Buffer A–100% Buffer B, 67.6min–95% Buffer A–5% Buffer B. Approximately 40–50 1-min fractions, representing peak peptide elution, were collected starting from initial peptides elution, and were reduced to dryness by vacuum centrifugation shortly thereafter. For downstream MS analysis, the fractions were concatenated into 20 samples by combining pairs of fractions which eluted at different time points during the gradient.

Each sample was analysed in an Orbitrap Eclipse mass spectrometer (Thermo Fisher Scientific). Mass spectra were acquired in positive ion mode applying data acquisition using synchronous precursor selection MS3 (SPS-MS3) acquisition mode^46^ triggered using Real-time Search against human protein sequences from UniProt/Swiss-Prot. Carbamidomethylation of cysteine and TMT-6plex (total proteome samples) or TMTpro-16plex tagging (Subcellular fractionated samples) of lysine and peptide N terminus were set as static modifications, with oxidation of methionine as a variable modification. Scoring thresholds were set as follows: Xcorr=1.4, dCn=0.1 and Precursor PPM=10. For details of LC-MS/MS acquisition and proteomics data processing see supplementary methods.

### MS spectra processing and peptide and protein identification

Raw data were viewed in Xcalibur v.2.1 (Thermo Fisher Scientific), and data processing was performed using Proteome Discoverer v2.3 (Thermo Fisher Scientific). The raw files were submitted to a database search using Proteome Discoverer with SequestHF and MS Amanda^47^ algorithms against the Homo sapiens database downloaded in June 2020 from UniProt/Swiss-Prot. Common contaminant proteins (several types of human keratins, BSA and porcine trypsin) from the common Repository of Adventitious Proteins (cRAP) v1.0 (48 sequences, adapted from the Global Proteome Machine repository, https://www.thegpm.org/crap/) were added to the database. The spectra identification was performed with the following parameters: MS accuracy, 10 p.p.m.; MS/MS accuracy of 0.5 Da; up to two missed cleavage sites allowed; carbamidomethylation of cysteine and TMT-6plex (total proteome samples) or TMTpro-16plex tagging (Subcellular fractionated samples) of lysine and peptide N terminus as a fixed modification; and oxidation of methionine and deamidated asparagine and glutamine as variable modifications. Percolator was used for false discovery rate estimation and only rank 1 peptide identifications of high confidence (FDR < 1 %) were accepted. TMT reporter values were assessed through Proteome Discoverer v2.3 using the Most Confident Centroid method for peak integration and integration tolerance of 20 p.p.m. Reporter ion intensities were adjusted to correct for the isotopic impurities of the different TMT reagents (manufacturer specifications). Reporter ion intensities were adjusted to correct for the isotopic impurities of the different TMT reagents (following manufacturer specifications). Sample labels for each TMT tag are presented in supplementary table 4.

### Spatial proteomics

Previous marker proteins defined for U-2 OS were annotated in^13^. 12 / 733 markers were deemed outliers based on manual curation of their profile and consideration of localisation assigned in HPA^48^ and GO. Nuclear proteins can diffuse out of the nucleus, complicating their abundance profile (see supplementary methods). To ensure accurate assignment of nuclear proteins, nuclear markers were annotated de novo, utilising the COMPARTMENTS database^49^. COMPARTMENTS localisations were mapped to the localisations defined by the standard LOPIT marker sets. Proteins with a score of 5 for the ‘Nucleus’ and no score over 2 for any other localisation were denoted as exclusively nuclear. The profiles for these nuclear exclusive proteins were split into three groups, which were separated based on the following thresholds on the row-sum normalised abundances: Group 1 =over 0.3 in pooled fraction 4. Group 2 = over 0.2 in pooled fraction 5. Group 3 = over 0.4 in pooled fraction 8. A GO enrichment analysis, using a Hypergeometric test, was then used to determine the enriched functionalities, relative to the background of all quantified proteins. This indicated that the first group of nuclear proteins were highly enriched in nucleolus, chromatin and ribosome biogenesis proteins, whereas the other two were enriched in nucleoplasm proteins but no more specific GO terms. Furthermore, proteins annotated as nuclear membrane or nuclear lamina in GO were more closely associated with group 1. Thus, group 1 was denoted as ‘Nucleus’ and the other groups were referred to as ‘Nucleoplasm-1’ and ‘Nucleoplasm-2’. Finally, the ‘Proteosome’ marker set was expanded to define a set of markers for ‘Protein complexes’ by including proteins annotated in GO as part of the MARS complex (‘GO:0017101’), COP9 signalosome (‘GO:0008180’) or eIF3 complex (‘‘GO:0005852’), with 5/42 of the additional protein complex markers excluded as outliers. SVM classification in basal conditions was performed as described in^50^, with hyperparameters selected by grid search and 50 iterations.

BANDLE^51^ was used to identify differentially localised proteins between control condition and UPR. Differential localisation analysis was performed on each replicate separately, with 10,000 MCMC iterations, 5,000 burn-in iterations, 4 chains and 1/20 thinning. Off diagonal values for the matrix of Dirichlet priors were set at 0.01, with default values used for the penalised complexity priors. MCMC chains were inspected for convergence based on the reported number of localisation outliers, as suggested in^51^, and all chains were found to converge. Localisations in each condition were determined by setting the following threshold: bandle localisation probability * (1 - bandle outlier probability) > 0.95. Where this threshold was not met, the localisation was deemed ‘Undefined’. Proteins were deemed differentially localised if they were never assigned to the same localisation across the two conditions and at least 2/3 replicates had BANDLE differential localisation probabilities over the following thresholds to define three levels of confidence were for relocalisation: >= 0.99 = ‘Highly confident’, >= 0.95 = ‘Confident’ and >= 0.85 = ‘Candidate’.

### RNA-seq data processing for CORALL RNA-Seq samples

RNA-Seq fastq processing and quantification was performed using bespoke pipelines built with CGAT-core^52^. Fastq files were demultiplexed using idemux (https://github.com/Lexogen-Tools/idemux) and concatenated into one fastq per sample. To assess PCR duplication rate, UMIs were extracted from the read sequences using UMI-tools ^53^. Reads were aligned to a concatenation of the hg38 reference genome and the artificial SIRV genome using hisat v2.2.1^54^ using default settings. Secondary reads and reads with MAPQ<10 were discarded. Duplicate reads were identified with UMI-tools dedup using default settings. Transcript isoform quantification was performed from the fastqs without deduplication, using Salmon v1.4.0^55^ against a concatenation of the ensembl v102 human transcriptome and the artificial SIRV annotations, using default settings.

### Data analysis

Data analyses post RNA and peptide quantification steps above were performed using R v4.0.3^56^ and R markdown notebooks^57^, making extensive use of the tidyverse R packages^58^, MSnbase^59^, pRoloc^60^ and camprotR (https://github.com/CambridgeCentreForProteomics/camprotR).

### Differential abundance analysis

Gene-level quantifications (transcripts per million; TPM) were parsed using tximport^61^. Differential gene abundance was tested using DESeq2^62^ with default settings, with a False Discovery Rate threshold of 5% used to identify significant changes in abundance.

### Spatial transcriptomics

Transcript isoform and gene-level quantifications (transcripts per million; TPM) were parsed using tximport^61^ to generate objects to hold the quantification estimates for a single experiment. To take advantage of the spatial proteomics functionalities available through pRoloc^60^, we stored the RNA quantification data in MSnSets. Separate objects were created to hold transcript isoform and gene-level abundance estimates. Transcripts and genes with average TPM < 0.5 or TPM==0 in >= 2/3 samples in a given condition were discarded. Post TPM-filtering, 41547 transcript isoforms and 14203 genes were quantified in at least 2 replicates in both conditions.

Localisation markers were identified using a combination of *a priori* markers and semi-supervised clustering by non-negative matrix factorisation (NNMF). *A priori* markers were defined as follows: Nuclear markers were > 16-fold nuclear enriched in nuclear/cytosol fraction RNA-Seq^63^ or manually defined known nuclear lncRNAs (XIST, MALAT1, MEG3, DLX6-AS1, PINCR, UCHL1-AS1, NEAT1). Cytosol markers were significantly enriched in NES APEX-Seq^5^, or manually defined from known cytosolic lncRNAs (LINCMD1, NORAD, H19, NKILA, SNHG5, DANCR, OIP5-AS1, SNHG1). ER markers were defined as having enrichment in ER-Ribo-Seq > 2^0.5 ^24^ and significant enrichment > 8-fold in KDEL APEX-Seq^5^ and a predicted signal peptide or transmembrane domain according to ensembl. Mitochondrial markers were mitochondrially-encoded mRNAs. To determine the optimal number of clusters (k) for NNMF, the imputation-based approach was used^64^, with the selected k value minimising the mean squared error for NNMF-imputation of randomly added missing values. This gave k=5 for the basal condition experiment and k=4 for the UPR experiment. NNMF cluster assignments in basal conditions were then compared to the a priori markers to define a set of NNMF-guided gene-level markers, in which each marker set was associated with the NNMF cluster with greatest overlap and all markers in the NNMF cluster were retained. In addition, the NNMF cluster containing the nuclear markers was further used to define a nucleolus marker set by intersecting it with the top 30 most abundant snoRNAs according to the total RNA-Seq. Finally, a novel NNMF cluster was observed in both basal condition and UPR, which contained few a priori markers. The genes assigned to the novel NNMF cluster in both basal and UPR conditions were used to define a novel profile, which was observed to have the greatest relative abundance in between the nucleus and cytosol profile peaks, and was hence denoted as cytosol-light. Gene-level markers were used to generate transcript-level markers by taking all the transcript isoforms for each gene-level marker where the ensembl transcript biotype and gene biotype matched. The final marker sets were manually curated to remove 14 / 135 gene-level markers and 37 / 202 transcript-level markers that were deemed to be outliers. The higher proportion of transcript-level markers removed reflects the markers having been built at the gene level and extended naively to transcript-level markers.

### Estimation of localisation proportions

RNA content per fraction was estimated using the relative abundance of all SIRV features and human RNA and the proportion of the fraction used for RNA-Seq library preparation. Relative RNA-Seq quantifications in each fraction were adjusted with respect to the RNA content per fraction and abundance estimates row-sum normalised across the 8 fractions per replicate. Nonnegative least squares regression was used to estimate localisation proportions by separately modelling the profile of each transcript/gene as a non-negative linear combination of the average profile for the markers of each localisation. Since most of the ER markers relocalised upon UPR and the mitochondrial markers represent 100% membrane localisation in both conditions (no RNA copies in the cytosol), mitochondrial markers were used to estimate the membrane proportion. The proportion estimates were bootstrapped 100 times by re-sampling the markers, with resampling. Proportion estimates for a given transcript/gene were discarded where the model did not account for at least 90% of the variance or the absolute value or the intercept was greater than 0.05. The mean proportions across replicates were estimated for transcripts/genes with estimates from at least 2/3 replicates. In total, proportions for 27368 transcript isoforms and 13274 genes were retained.

Reads were frequently observed to bridge annotations between lncRNAs and neighbouring protein-coding, which resulted in misestimation of lncRNA proportions. To avoid this, we excluded all lncRNAs with a protein-coding gene within 15Kb downstream or 30Kb upstream.

### Defining signal peptide and transmembrane domain features

Signal peptide and transmembrane (TM) domain annotations were obtained from ensembl v102 and the distance between the start of the first signal peptide or TM domain and the stop codon was determined. For gene-level analyses, the minimum distance across all transcript isoforms was used. A conservative annotation of presence/absence of either signal peptide and/or TM domain was obtained by taking the union of the ensembl annotations with uniprot annotations.

### eCLIP binding data

eCLIP data in bed format was obtained from ENCODE using ENCODExplorer. Genomic coordinates were converted to transcriptomic coordinates using mapToTranscripts function in GenomicFeatures R package^65^.

### Modelling lncRNA cytosol localisation

To model the cytosolic localisation of lncRNAs, transcript length, spliced status, eCLIP binding data, kmers frequencies, AU content, RNA modifications, PolyA status, presence in FANTOM5 robust catalogue, and abundance were considered. Transcript length and splicing status was obtained from ensembl v102. eCLIP data was obtained as described above. 64 RBPs with > 10 lncRNA targets were retained and binding data converted to binary 0=unbound, 1 =bound. Kmers (k=1-7) were counted and expressed as frequencies. RNA modifications were obtained from m6A-atlas^66^, with modifications with > 10 lncRNA targets retained, namely, m6A, m5c, m1A and Psi. The FANTOM5 robust catalogue and PolyA status were obtained from^67^. PolyA status was converted to a binary 0=non-polyadenylated, 1=polyadenylated, with ‘undetermined’ encoded as 0 and ‘bimorphic’ encoded as 1. Abundance was calculated as the mean TPM from the total RNA-Seq of basal condition samples. Cytosol proportions were converted to a binary feature, where 1 = cytosolic (⅔ cytosol) and 0 = not-cytosolic (< ⅓ cytosol). The data was then split 80:20 into training and test data. Elastic net logistic regression was used to model the cytosol variable, with 10-fold cross-validation, using the glmnet R package^68^. By default, glmnet scales the dependent variables but returns the coefficients on the original scale. Alpha values were varied between 0-1, in steps of 0.1. Cross-validation folds were pre-computed to ensure they were the same for each alpha value. The alpha value with the minimum mean cross validation error was identified as 1, e.g lasso regression. Lambda was selected to give the most parsimonious model, using the ‘one-standard-error’ rule^69^. The final model contained 45 / 21918 non-zero coefficients. The predictive accuracy of this model was assessed using the hold-out test data and compared to the accuracy of a simpler logistic regression model of just transcript length, polyA status and AU content.

### Modelling UPR-resistant membrane localisation

Membrane proportion in UPR was modelled as being dependent upon membrane proportion in control conditions using a Generalised Additive Model (GAM) with a cubic regression spline with shrinkage, using the mgcv R package. The residual from the GAM was taken to represent the degree of UPR resistant localisation relative to the overall reduction in membrane proportions. Lasso regression (glmnet R package^68^) was used to model UPR resistance, with RBP binding from eCLIP data (see above) being the dependent variables. 10-fold cross-validation was used to select the lambda value that minimised the mean cross validation error, with the ‘one-standarderror’ rule used to select the most parsimonious model^69^. Transcripts with signal peptides/TM domains were separately modelled transcripts from those without.

### Modelling relocalisation to granules

Granule relocalisation (UPR granule proportion - Control granule proportion) was modelled according to the following features which were annotated to each mRNA: localisation in basal conditions, RNA length, RBP binding, sequence kmers, 5’ AUGs, codon features, upstream open reading frames (uORFs) and internal ribosome entry site (IRES). One-hot encoding was used for the features describing the localisation in basal conditions, where the highest proportion localisation was taken to be the single localisation for the RNA. Transcript, 5’ UTR, 3’UTR and coding sequence lengths were obtained from ensembl v102. eCLIP data was obtained as described above. 177 RBPs with > 100 mRNA transcript targets were retained and binding data converted to binary 0=unbound, 1=bound. Kmers (k=1-6) were counted and expressed as frequencies. 5’ AUGs were identified with separate features to encode in-frame and out-of frame AUGs. Codon, dinucleotide and wobble base frequencies were computed from the coding sequence. Codon optimality (MILC^70^) was computed, with separate features for the comparison against all transcripts and just the top 1% most abundant. uORFs were identified from^71^, retaining only ORFs with a score > 5 and an AUG start codon. Annotated IRES were obtained from IRESbase^72^. uORF and IRES were converted to binary presence/absence features for each transcript. In addition, pairwise interactions between the localisation in basal conditions and the transcript, 3’UTR, 5’UTR and coding length features and RBP binding features were added as separate features.

Granule relocalisation data was split 80:20 into training and test data. Elastic net regression was used to model the granule relocalisation, with 10-fold cross-validation, using the glmnet R package^68^. By default, glmnet scales the dependent variables but returns the coefficients on the original scale. Alpha values were varied between 0-1, in steps of 0.1 Cross-validation folds were pre-computed to ensure they were the same for each alpha value. The alpha value with the minimum mean cross validation error was identified 1, e.g lasso regression. Lambda was selected to give the most parsimonious model, using the ‘one-standard-error’ rule^69^. The final model contained 74 / 16895 non-zero coefficients.

### Data processing and analysis for differential centrifugation-based LoRNA

Data processing and analysis for samples quantified using 3’ Quant Seq was identical to CORALL RNA-Seq samples, except for the following: Gene read counts were normalised to counts per million (CPM) using the total number of assigned reads per sample. Genes with average CPM < 1 or CPM==0 in > 20% of the samples were discarded. Markers identified from equilibrium density centrifugation-based LoRNA were annotated and proportions calculated as indicated above, except that proportions were estimated separately for mitochondria and ER and summed to give membrane proportions, and ‘cytosol-light’ proportions were not estimated. The mean proportions across replicates were estimated for transcripts/genes with estimates from at least 3/5 replicates. For details on the analysis of the technical bias between RNA length and sedimentation, see supplementary methods.

### Gene-Ontology enrichment analyses

All Gene-Ontology (GO) enrichment analyses were performed using the goseq R package^73^. For the analysis of GO terms enriched in eIF3d-bound membrane RNAs, the membrane proportion in basal conditions was included as the bias factor and enrichment effect sizes accounting for biasing factors were calculated using the probability weight functions obtained with goseq using the estimate_go_overrep function in camprotR. For the analysis of GO terms enriched in the membrane RNAs not encoding a signal peptide or TM domain, no bias factor was included and a Hypergeometric test was instead used. For both analyses, p-values were adjusted for multiple testing using the Benjamini-Hochberg procedure^74^ and GO terms with adjusted p-value > 0.05 (5% FDR) or accounting for fewer than 10% of the foreground genes were excluded. Redundant GO terms were removed using the remove_redunant_go function in camprotR R package.

### Single molecule FISH probe design and synthesis

Subcellular RNA localisation was assessed by using an adaptation of the single-molecule inexpensive FISH protocol^75^. Z (CTTATAGGGCATGGATGCTAGAAGCTGG) and Y (AATGCATGTCGACGAGGTCCGAGTGTAA) FLAP DNA handles, labelled at the 5’ and 3’ ends with Atto488, were purchased from Sigma (HPLC-purified) and resuspended to a concentration of 100 μM in nuclease-free water. For each target RNA, 30-48 DNA probes of 20 nucleotides (nt) were designed with a minimum spacing length of 2 nt and a guanine-cytosine content of 40-65%. Each gene-specific sequence was flanked by a 28 nt sequence complementary to either a Z- or Y-FLAP sequence. The resulting 48 nt probes were purchased from Sigma (standard desalt purification, 100 μM in nuclease-free water). Fluorescently labelled gene-specific probes were then generated as follows: 200 pmol of an equimolar mixture of all gene-specific oligos for each gene were mixed with 250 pmol of the appropriate FLAP oligo in 1x NEBuffer 3 (New England Biolabs, B7003), then incubated in a Thermocycler (BioRad) for 3 min at 85 °C, 3 min at 65 °C, and 5 min at 25 °C (lid 99°C).

### smiFISH/Immunofluorescence and Immunofluorescence

For smiFISH/IF experiments, 8 104 U 2-OS cells were seeded in each well of a 12-well plate, on top of no 1.5 glass coverslips previously washed in 1 M HCl. The following day, cells were treated with either 250 nM Tg or the corresponding volume of DMSO, rinsed three times with PBS (with MgCl_2_ and CaCl_2_, Sigma D8662), then fixed for 10 min in 3% Methanol-free PFA (Alfa Aesar, 43368) in PBS at RT. The fixative was then quenched in 100 mM Glycine (Sigma) in PBS for 10 min at RT, samples were then washed twice in PBS for 10 min and permeabilised in 70% EtOH at 4 °C for at least 1 h. Samples were then prepared for hybridisation by washing in 10% Formamide (Sigma), 1 U/μl RNasin Plus (Promega) in 2x Saline-sodium citrate (SSC) buffer for 10 min. From this step onwards, samples were protected from direct light. Hybridisation was performed by incubating coverslips with 100 - 250 nM probes diluted in hybridisation buffer (2X SSC buffer, 10% Dextran sulfate (Sigma), 10% Formamide, 2 mM ribonucleoside vanadyl complexes (Sigma), 200 μg/ml bovine serum albumin (Roche), 1 mg/ml E. coli tRNA (Roche), 1 U/μl RNasin Plus (Promega)), for 3 h at 37 °C in a humid chamber. Coverslips were then transferred to a clean 12-well plate and washed twice in 10% Formamide, 1 U/μl RNasin Plus, 2x SSC buffer, for 10 min at 37 °C. Further washes were then performed (three times in 2X SSC with no incubation, twice in PBS for 10 min), before incubation with blocking buffer (3% nuclease-free bovine serum albumin (Sigma) in PBS) for 30 min at RT. Samples were then incubated with primary antibody diluted 1:500 in blocking buffer, for 2 h at RT. They were then washed three times in PBS for ten min, incubated with a secondary antibody diluted 1:2000 in blocking buffer, for 1 h at RT. After three additional washes in PBS, the nucleus was stained by incubation with 4’,6-diamidino-2-phentylindole (DAPI, Sigma, 200 ng/ml in PBS) for 1 min at RT. Coverslips were then washed twice in PBS for 5 min before being mounted onto glass microscope slides with a drop of ProLong™ Glass Antifade Mountant (Invitrogen).

For consistency, IF experiments were performed following the smiFISH/IF procedure with the omission of the FISH hybridisation step and the washes in 10% formamide, 2X SSC.

### Confocal microscopy

Images were acquired on a Zeiss Axio Observer.Z1 LSM 980 microscope equipped with Airyscan 2, using ZEN Blue software (version 3.3), in Airyscan super-resolution imaging mode. A C-Plan-Apochromat 63X /1.4 NA oil objective was used with Zeiss Immersol 518F (23 °C) immersion oil. 20 Z-slices per image were acquired at an interval of 0.13 μm.

### Image processing and quantification

Airyscan processing with standard parameters was applied to each image in Zen Blue software. Further processing and analysis were performed using FiJi software^76^ (version 2.3.051). RNA FISH images were background subtracted (rolling ball radius = 10 pixels); RNA foci were then detected and counted using the FindFoci function, as part of the GDSC plugin^77^. To quantify the degree of co-localisation of RNA foci with the ER, Manders’ overlap coefficient was calculated using the JACoP plugin^78^.

### Cell migration assay

Cell migration assays were performed using the xCELLigence RTCA DP instrument (ACEA Biosciences) according to manufacturer’s instructions. Cells were treated with thapsigargin for 1 hour, collected by trypsinisation and washed in PBS to remove all traces of serum. 30,000 cells were seeded in the upper chamber of a 16 well migration plate (CIM-16 plate) in 100 μl of media containing 0.1% serum. Cells migrate to a lower chamber containing 160 μl of media supplemented with 10% serum. As cells migrate across the microelectrodes into the lower chamber they generate impedance measurements which enables label free quantification of cell migration for 24 hours (Cell index). Cell indexes for Tg-treated samples were expressed as fold changes relative to the DMSO control sample. Significance testing between eIF3d and control siRNA knockdowns was performed with a paired Student’s t-test. For details on cell death assays, see supplementary methods.

## Supporting information

Supplementary Table 1 - LoRNA_normalised_quantifications

Supplementary Table 2 - LoRNA_proportions

Supplementary Table 3 - LOPIT

Supplementary Table 4 - Experimental_details

## Data availability

The mass spectrometry proteomics data have been deposited to the ProteomeXchange Consortium via the PRIDE^79^ partner repository with the dataset identifier PXD030456. The RNA-Seq data have been deposited in ENA with study accession PRJEB49479.

## Acknowledgements

We would like to thank Dee Scadden for kindly providing the G3BP1-GPF expression plasmid, B. Fisher for kindly sharing equipment and Maria Marti-Solano for assistance with the manuscript writing. E.V., T.S., R.M.L.Q., R.F.H., M.P. and ME are supported by Wellcome Trust, grant numbers 110170/Z/15/Z and 110071/Z/15/Z awarded to A.E.W. and K.S.L. V.D. is supported by Medical Research Council, grant number 5TR00. LMB is supported by EU Horizon 2020 programme INFRAIA project EPIC-XS(Project 823839). OMC was supported by a Wellcome Trust Mathematical Genomics and Medicine student funded by the Cambridge School of Clinical Medicine and by the Todd-Bird Junior Research Fellowship from New College, Oxford.

## Author contributions

**Eneko Villanueva**:Conceptualization, Methodology, Investigation, Writing - Original Draft, Writing - Review & Editing, Visualization, Project administration. **Tom Smith**: Conceptualization, Methodology, Formal analysis, Investigation, Data Curation, Writing - Original Draft, Writing - Review & Editing, Visualization, Project administration. **Mariavittoria Pizzinga**: Methodology, Formal analysis, Investigation, Visualization, Writing - Original Draft, Writing - Review & Editing, Visualization. **Mohammed Elzek**: Methodology, Investigation, Writing - Review & Editing. **Rayner M. L. Queiroz**: Investigation, Writing - Review & Editing. **Robert F. Harvey**: Investigation, Writing - Review & Editing. **Lisa M Breckels**: Resources. **Oliver M. Crook**: Formal analysis, Writing - Review & Editing. **Mie Monti**: Writing - Review & Editing. **Veronica Dezi**: Writing - Review & Editing. **Anne E. Willis**: Funding acquisition, Conceptualization, Project administration, Supervision, Writing - Review & Editing. **Kathryn S. Lilley**: Funding acquisition, Conceptualization, Project administration, Supervision, Writing - Review & Editing

