## Supplementary Methods for "A system-wide quantitative map of RNA and protein subcellular localisation dynamics"

LC-MS/MS acquisition

TMT-labelled samples were analysed in an Orbitrap Eclipse coupled to a nanoLC Dionex Ultimate 3000 UHPLC (Thermo Fisher Scientific). Peptides were trapped on a 100 µm × 2 cm, C18, 5 µm, 100 trapping column (Acclaim PepMap 100) in µl-pickup injection mode at 15 µl/min flow rate for 3 min. Samples were then loaded on a Rapid Separation Liquid Chromatography, 75 µm × 50 cm nanoViper C18 3 µm 100 column (Acclaim, PepMap) at 50 ºC retrofitted to an EASY-Spray source with a flow rate of 300 nl/min. Analytical chromatography was performed over 120 min (buffer A, HPLC $H_2O$, 0.1% formic acid; buffer B, 100% ACN, 0.1% formic acid; 0–3 min: at 2% buffer B, 3–105 min: linear gradient 2% to 40% buffer B, 105-105.3 min: 40% to 90% buffer B, 105.3-110 min: at 90% buffer B, 110-110.3 min: 90% to 3% buffer B, 100.3-120 min: at 3% buffer B). Each MS1 scan was performed in the Orbitrap analyser (mass range = m/z 400–1500, resolution = 120,000). Precursors with charge between 2 and 6 and intensity above 5000 were selected for collision induced dissociation (CID)-MS2 fragmentation, with normalised AGC target of 200% and maximum accumulation time of 50 ms. Mass filtering was performed by the quadrupole with 0.7 m/z transmission window, followed by CID fragmentation in the linear ion trap with 30% normalized collision energy. Selected fragmented ions were dynamically excluded for 60 s. Real-time Search (RTS) was used to trigger SPS-MS3 acquisition. RTS used Human UniProt/Swiss-Prot database, carbamidomethylation of cysteine and TMT-6plex (total proteome samples) or TMTpro-16plex tagging (Subcellular fractionated samples) of lysine and peptide N terminus as static modification, oxidation of methionine as variable modification, as scoring thresholds Xcorr=1.4, dCn=0.1, Precursor PPM=10, one missed cleavage and maximum search time of 35 ms. SPS was applied to co-select 10 fragment ions for HCD-MS3 analysis. SPS ions were all selected within the 400–1,500 m/z range and were set to preclude selection of the precursor ion and TMT or TMTpro ion series. Normalised AGC targets and maximum accumulation times were set to 200% and 120 ms. Co-selected precursors for SPS-MS3 underwent HCD fragmentation with 55% normalised collision energy and were analysed in the Orbitrap with nominal resolution of 50,000. The number of SPS-MS3 spectra acquired between full scans was restricted to a duty cycle of 3 s.

Proteomics data processing

PSM-level quantifications were filtered to conservatively remove peptides from common contaminants. Alongside the cRAP proteins, further potential contaminants were identified by considering all proteins that shared an observed peptide with a cRAP protein as further contaminants. PSMs without a unique master protein assigned or more than 20% missing values were excluded. Remaining missing values were imputed using knn imputed (k=10), with sum normalisation prior to imputation and de-normalisation post imputation, to ensure nearest neighbours shared similar abundance profiles over the fractions, rather than similar average abundance. We then identified and removed outlier PSMs which had a median euclidean distance over 0.2 from all other PSMs for the same master protein. In doing so, PSMs with higher co-isolation and lower average signal/noise were selectively removed where these low quality PSMs disagreed with other PSMs. Remaining PSMs were then median-centre-normalised. Protein-level abundances were estimated by summing PSM-level abundances, for proteins with at least two PSMs. Protein abundances were row-sum normalised such that the total abundance across all fractions from a given replicate equaled one.

Assessing the technical bias for RNA length and sedimentation

Gene-level CeFra-Seq quantification data (TPM) was downloaded using the ENCODExplorer R package. We computed the gene length as the mean length of all transcript isoforms included in the GENCODE basic gene set and with Transcript Support Level 1 in ensembl. Mean abundances per fraction were computed for the cytosol and membrane fractions, from which ratios were then computed compared to the total RNA-Seq samples. High-confidence cytosolic and membrane localised RNAs were obtained from LoRNA, setting a threshold of proportion > 80%. Pearson product-moment correlation coefficients were computed for gene length vs cytosol/total for cytosolic RNAs and membrane/total for membrane RNAs. The same procedure was used for the differential sedimentation-based RNA mapping approach estimates, with supernatant/total correlated with gene length for cytosolic RNAs.

Cell death assay

All flow cytometry data was acquired using a BD LSRFortessa (BD bioscience) and analysed with BD FACSDiva software (version 9.0.1). 10,000 counts were acquired for each experimental condition. Cells were collected in Annexin binding buffer (BD biosciences #556454) and cell death was determined after incubation with Annexin-V-FITC (Thermo Fisher, BMS500FI-100) and Draq7 (Abcam, ab 109202).

Polysome profiling

10%-50% (w/v) sucrose gradients were prepared in gradient buffer (100 mM NaCl, 5 mM $MgCl_2$, 15 mM tris-HCL pH 7.5, 1 mM DTT, 0.1 mg/ml cycloheximide). Cells were washed in PBS-cyclohexamide (100 g/ml) and scraped into lysis buffer (100 mM NaCl, 5 mM $MgCl_2$, 15 mM tris-HCL pH 7.5, 1 mM DTT, 0.2 M sucrose, 0.1 mg/ml cycloheximide, 0.5% IGEPAL, 5 µl RNasin per 1 ml). Lysates were incubated on ice for three min and cells pelleted by centrifugation at 1300 x g for 5 min. The supernatant was layered on top of a gradient and centrifuged at 38,000 rpm (acceleration 9, deceleration 6) for 2 h at 4 $^{o}$C using a Beckman Coulter ultracentrifuge. Polysome profiles were obtained by measuring absorbance at 254 nm using a UA-6 UV-VIS detector (Presearch Ltd).

Reporter gene expression analysis

DCP2-GFP tagged mammalian cell expression plasmid was directly acquired from Adgene (pT7-EGFP-C1-HsDCP2, ref 25031). G3BP1-GFP plasmid was kindly provided by Dr. Dee Scadden (University of Cambridge). Plasmids were purified using EasyPep™ Mini MS Sample Prep Kit (Quiagen). One plate of 500 $mm^2$ of 90% confluent U-2 OS cells were transfected per plasmid, per condition using 100 µg of plasmid with Lipofectamine 3000 Transfection Reagent (Thermo Scientific) according to manufacturer instructions. Two days post-transfection, cells were treated with DMSO or TG as indicated above and lysed and fractionated as specified in the "density based cell fractionation" methods section. Gradient fractions were imaged on a Zeiss Axio Observer.Z1 LSM 980 microscope and fluorescence intensity quantified with ImageJ 1.8.0_172 for Mac OS X.

Resolving nuclear proteins

While RNA export from the nucleus is a tightly regulated process, small proteins diffuse freely. Therefore, when interpreting cell fractionation experiments it is important to consider that small nuclear proteins with weak interactions leak from the nucleus[80]. While fixing protein-protein interactions could overcome this limitation, fixing agents like FA negatively affect organelle resolution making this approach currently incompatible with density gradient fractionation (Supplementary methods Fig 1.). Future advances in reagents to efficiently stabilise protein-protein interactions without affecting organelle integrity and lipid content may help overcome this limitation. Here, we selected nucleoplasm protein markers reflecting this phenomenon to avoid miss-classification of nucleoplasm proteins to cytosol, achieving a cytosolic marker F1 score of 0.85.

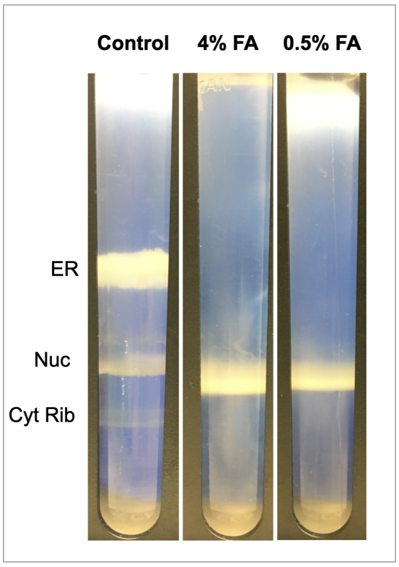


**Supplementary methods figure 1. Formaldehyde crosslinking compatibility with density centrifugation cell fractionation**
Representative images of the banding pattern of cell lysates after density fractionation in non-crosslinked and formaldehyde crosslinked conditions.
