## Supplementary Figures for "A system-wide quantitative map of RNA and protein subcellular localisation dynamics"

### Supplementary Figure 1.

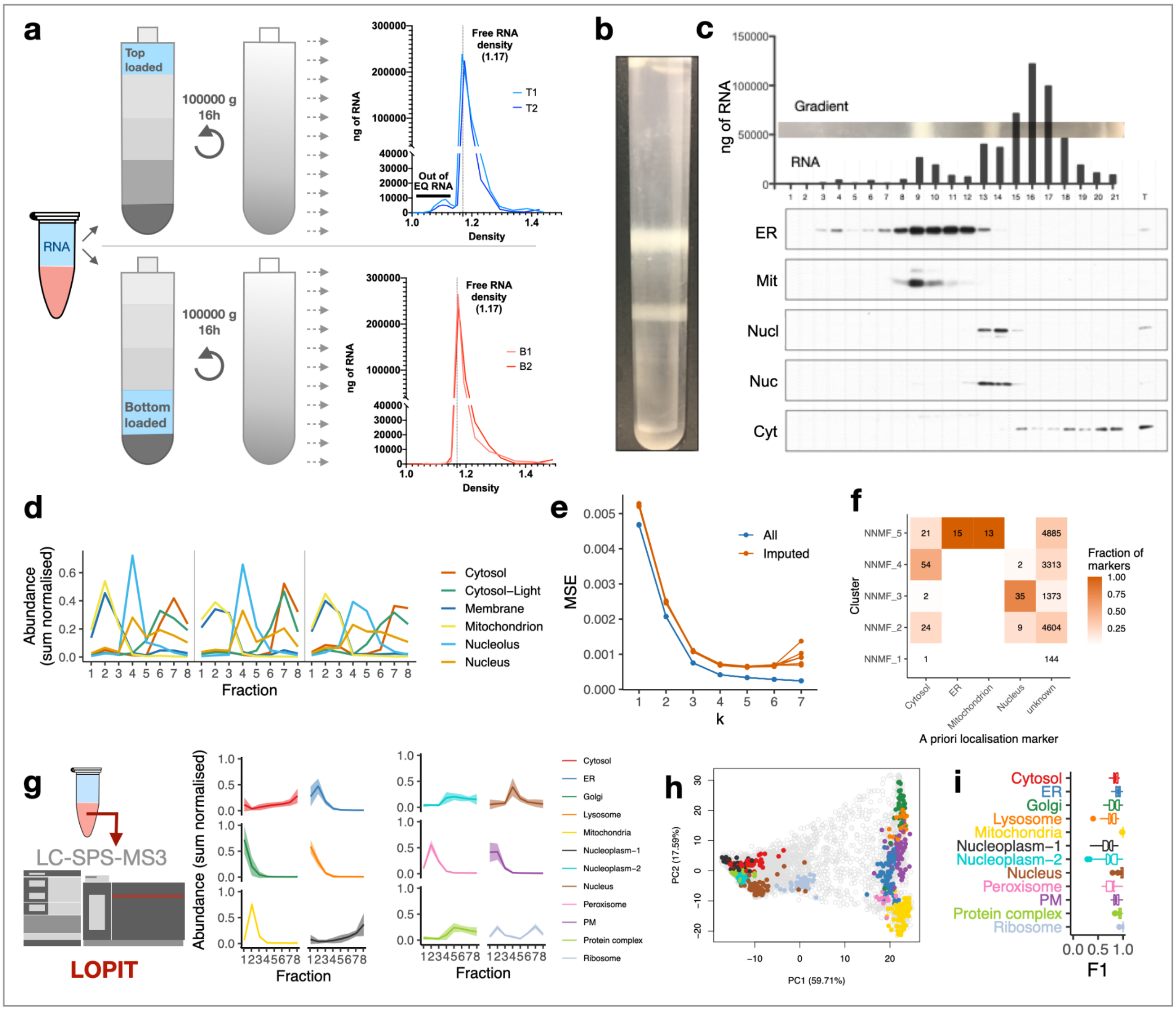


#### Simultaneous maps of RNA and protein subcellular localisation

(a) Cell-free RNA sedimenting profile along the density gradient. T = top loading (blue), B = bottom loading (red, at 1.17 g/ml). Fractions with density < 1.17 are enriched in organelles.
(b) Representative image of a density-based cell fractionation gradient post centrifugation.
(c) Quantification of RNA (in ng) at the different gradient fractions, and a representative western blot of ER (Calreticulin), mitochondria (Mit, Cytochrome c oxidase subunit 4), nucleolus (Nucl, Fibrilarin) , nucleus (Nuc, Histone H3) and cytosolic (b-Actin) compartment markers
(d) Linear profiles of the indicated RNA markers per replica
(e) Selection of optimal K for NNMF clustering of gene-level profiles in control condition. MSE = Mean Standard Error. Imputed = data points that were removed and imputed by NNMF. MSE lowest for imputed data points at k=5.
(f) Intersections between highest scoring NNMF cluster and a priori markers, where ‘unknown’ denotes non-markers.

(g) Protein quantification by LC-MS3 and representative protein marker profiles. Line and shaded region represents mean +/- one standard error.
(h) Kernel PCA of protein makers.
(i) Distributions of F1 scores for protein markers for each localisation using support vector machine classification. n = 50 iterations.

**Supplementary Figure 2.**

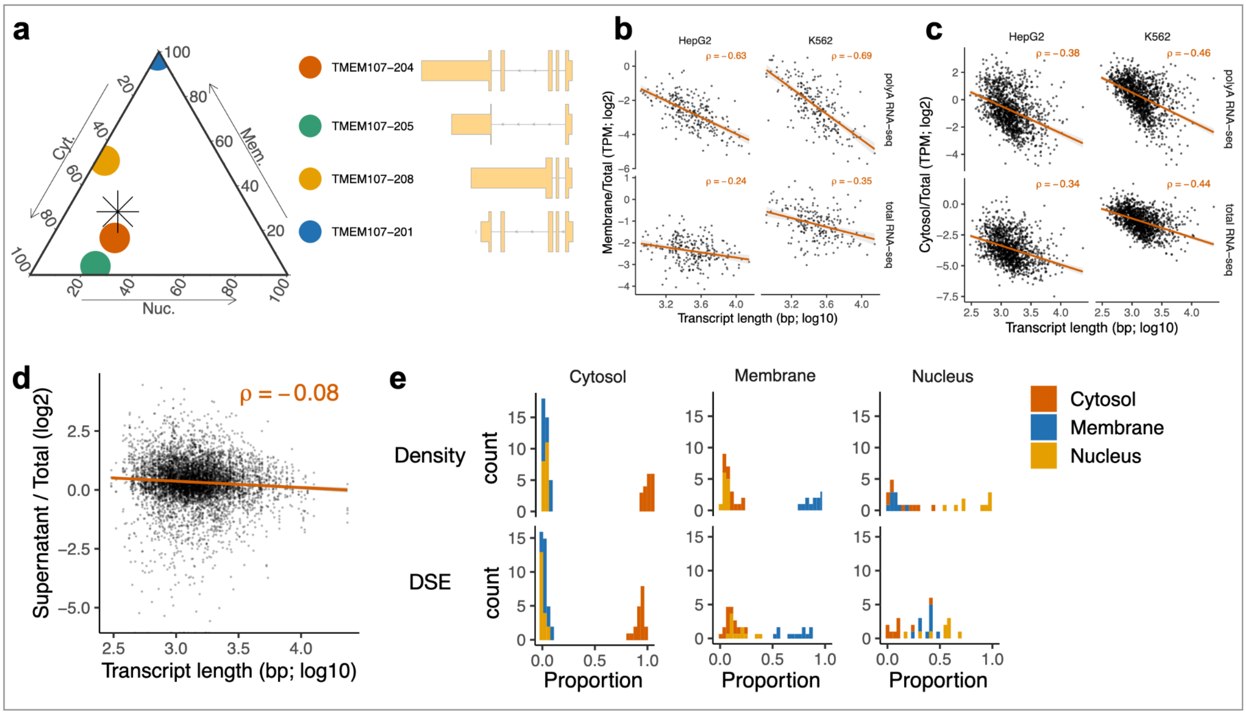


**System-wide quantification of RNA localisation**

(a) Cytosol, membrane and nucleus proportions for TMEM107 transcript isoforms. Gene-level proportion indicated with asterisk (left). Transcript models (right).
(b) Transcript length vs CeFra-Seq (ref) membrane / total for RNAs with > 80% membrane proportion according to equilibrium density centrifugation-based LoRNA.
(c) As per b, except cytosol / total for RNAs with > 80% cytosol proportion
(d) As per b, except differential centrifugation speed-based LoRNA final fraction / total for RNAs with > 80% cytosol proportion
(e) Proportions for cytosol, membrane and nucleus markers using equilibrium density centrifugation ('Density') and differential sedimentation ('DSE') based cell fractionation

**Supplementary Figure 3.**

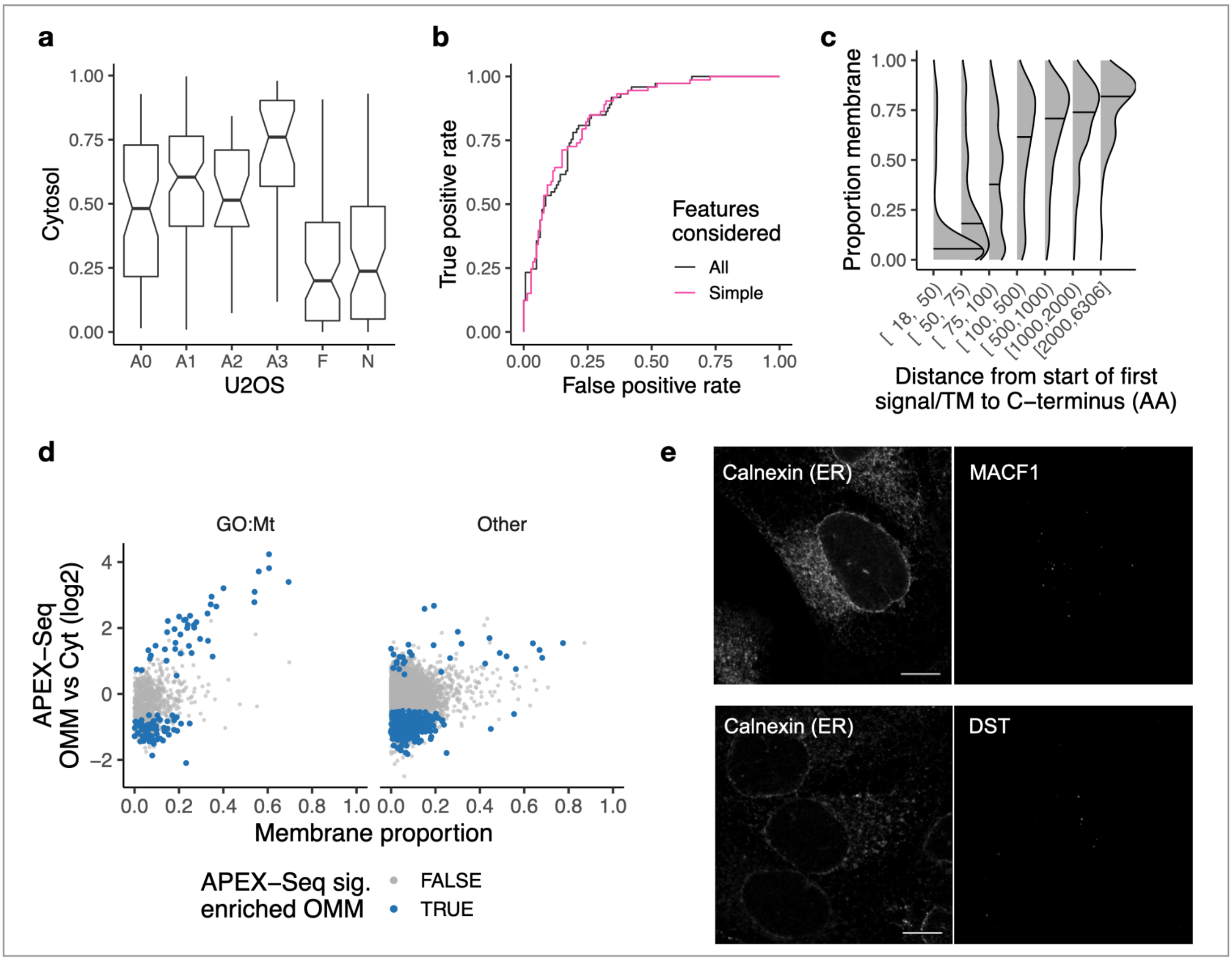


### Features driving RNA localisation

(a) Cytosol proportions for lncRNAs annotated by their ribosome association[22]. A0-A3 represent increasingly confident ribosome association. F=Not associated. N=Undetermined.
(b) Receiver operating characteristic curves for logistic regression models of lncRNA cytosol localisation. All=Lasso regression using all features. 'Simple'=Just using AU content, polyA status and transcript length.
(c) Relationship between the membrane proportion of an mRNA and the distance from the first signal peptide or transmembrane domain to the stop codon of the polypeptide product.
(d) RNAs annotated as encoding mitochondrial proteins in GO and also enriched in APEX-Seq OMM (vs cytosol) have high membrane proportions in LoRNA.
(e) Representative microscopy images showing smFISH for MCF1 (top-right) and DST (bottom-right) in parallel with the ER (left panels) immuno-labelled with an anti-calnexin antibody. Scale bar = 10 μm.

**Supplementary Figure 4.**

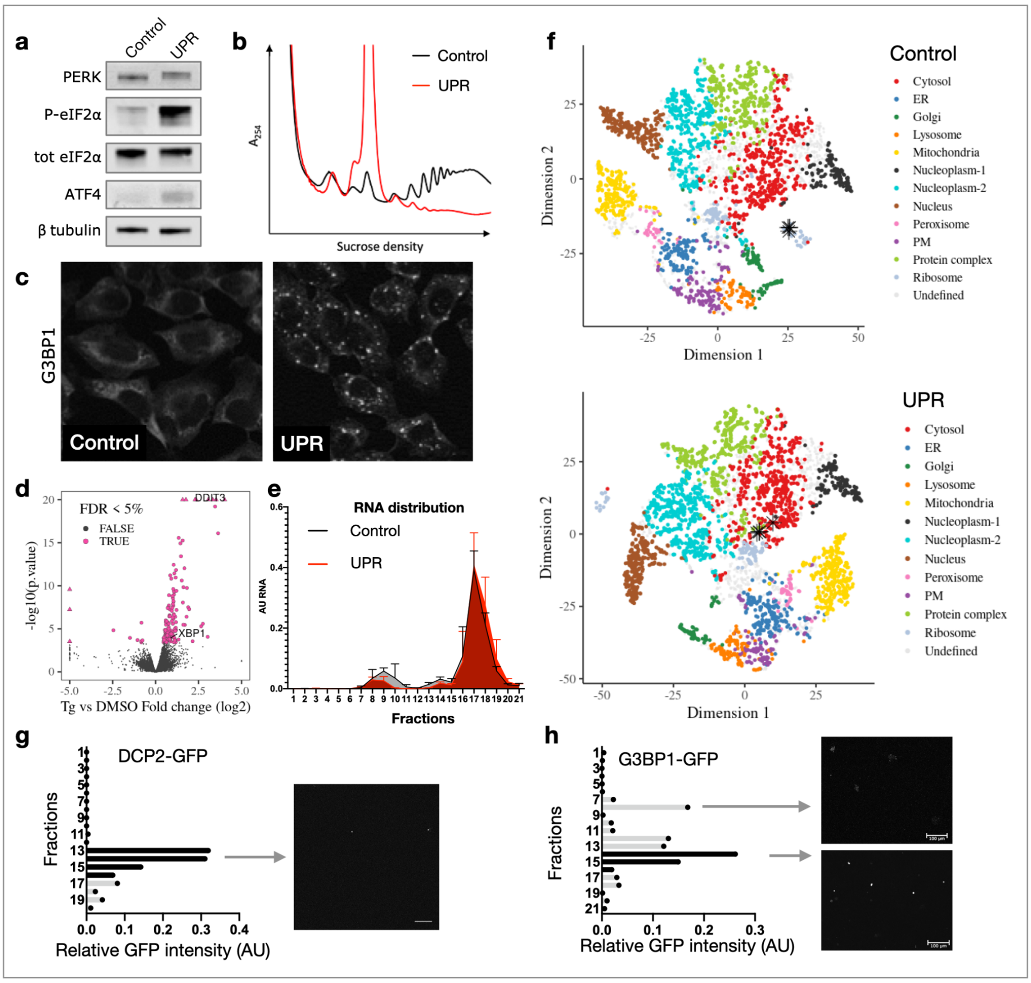


### Transcriptome and proteome subcellular redistribution upon UPR

(a) Representative western blot for samples treated with DMSO or TG showing canonical UPR markers. TG treatment induces: a shift in PERK size, consistent with phosphorylation; phosphorylation of eIF2α; increased expression of stress-specific transcription factor ATF4.
(b) Representative polysome profiles in control (DMSO) and UPR (1 h at 250 mM thapsigargin). For experimental details, see supplementary methods.
(c) Representative Z-projection images of control (DMSO) and UPR-induced (TG 1h) U-2 OS cells immuno-labelled with an antibody for stress granule marker G3BP1.
(d) RNA abundance changes between UPR and control. Significant changes (5% FDR) highlighted. DDIT3/CHOP and XBP1 labelled.
(e) RNA profile along unpooled density gradient fractions in control and UPR conditions. Fractions 1:11 represent light organelles (including mitochondria and ER), fractions 13 and 14 nucleus, and 15 to 21 cytosol (n = 3 per condition).
(f) t-SNE projections for protein localisation profiles in Control and UPR. The four proteins that relocalise away from the ribosome and towards the cytosol light profile are highlighted with asterisks. Proteins are coloured by their localisation allocation from BANDLE. Proteins with inconsistent localisation allocation are denoted as ‘Undefined’.
(g) DCP2-GFP sedimentation profile along unpooled density gradient fractions measured as GFP fluorescence intensity per fractions. Fractions corresponding to the density range of the cytosol light (pooled fraction 6 in Fig 1) in black (left), and representative images of the peaking fraction containing DCP2 granules respectively (right). Scale bar = 100 µm.

(h) G3BP1-GFP sedimentation profile along unpooled density gradient fractions under UPR stimulation measured as GFP fluorescence intensity per fractions. Fractions corresponding to the density range of the cytosol light in black (left), and representative images of fractions 7 and 14 representing membrane enclosed protein aggregates and G3BP1 containing granules respectively (right). Scale bar = 100 µm.

**Supplementary Figure 5.**

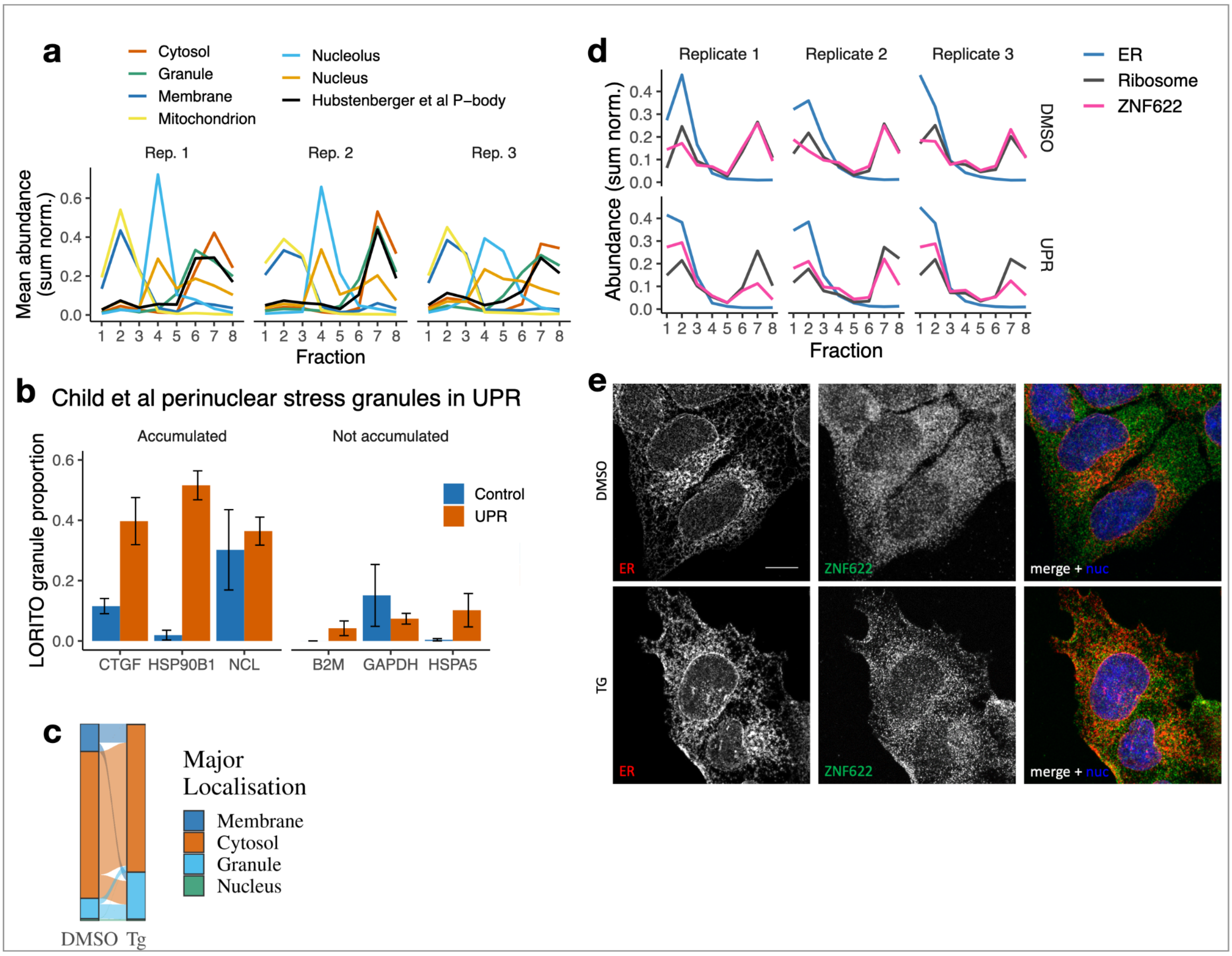


**Analysis of the characteristics driving RNAs to granules**

(a) Mean profiles for RNA markers across the 3 replicates in control conditions.
(b) Granule proportions for RNAs found to accumulate/not accumulate in stress granules upon UPR[34].
(c) Primary localisation of transcripts in control and UPR.
(d) Abundance profile of ZNF622 in Control and UPR, with average profiles for ER and ribosome markers.
(e) Representative Z-projections of control (DMSO) and UPR-induced (TG) U-2 OS cells immuno-labelled with antibodies for calnexin (ER marker) and ZNF622, and co-stained with DAPI. Scale bar = 10 µm.

**Supplementary Figure 6.**

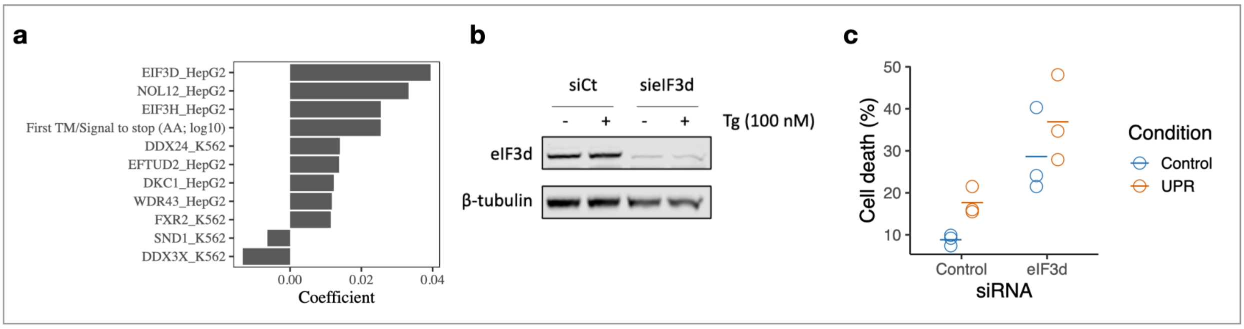


**Analysis of the RNAs retaining membrane association under UPR**

(a) Coefficients for RBPs which are predictors of GAM residuals for RNAs which encode a signal peptide/TM domain
(b) Western blot validating eIF3d knockdown.
(c) Quantification of cell death using Annexin V-FITC and Draq7 staining following eIF3d knockdown. Cells were treated with a 1 hour pulse of thapsigargin and allowed to recover for a further 24 hours before.
